# Neutralizing Gatad2a-Chd4-Mbd3 Axis within the NuRD Complex Facilitates Deterministic Induction of Naïve Pluripotency

**DOI:** 10.1101/192781

**Authors:** Nofar Mor, Yoach Rais, Shani Peles, Daoud Sheban, Alejandro Aguilera-Castrejon, Asaf Zviran, Dalia Elinger, Sergey Viukov, Shay Geula, Vladislav Krupalnik, Mirie Zerbib, Elad Chomsky, Lior Lasman, Tom Shani, Jonathan Bayerl, Ohad Gafni, Suhair Hanna, Jason D. Buenrostro, Tzachi Hagai, Hagit Masika, Yehudit Bergman, William J. Greenleaf, Miguel A. Esteban, Yishai Levin, Rada Massarwa, Yifat Merbl, Noa Novershtern, Jacob H. Hanna

## Abstract

The Nucleosome Remodeling and Deacytelase (NuRD) complex is a co-repressive complex involved in many pathological and physiological processes in the cell. Previous studies have identified one of its components, Mbd3, as a potent inhibitor for reprogramming of somatic cells to pluripotency. Following OSKM induction, early and partial depletion of Mbd3 protein followed by applying naïve ground-state pluripotency conditions, results in a highly efficient and near-deterministic generation of mouse iPS cells. Increasing evidence indicates that the NuRD complex assumes multiple mutually exclusive protein complexes, and it remains unclear whether the deterministic iPSC phenotype is the result of a specific NuRD sub complex. Since complete ablation of Mbd3 blocks somatic cell proliferation, here we aimed to identify alternative ways to block Mbd3-dependent NuRD activity by identifying additional functionally relevant components of the Mbd3/NuRD complex during early stages of reprogramming. We identified Gatad2a (also known as P66α), a relatively uncharacterized NuRD-specific subunit, whose complete deletion does not impact somatic cell proliferation, yet specifically disrupts Mbd3/NuRD repressive activity on the pluripotency circuit during both stem cell differentiation and reprogramming to pluripotency. Complete ablation of Gatad2a in somatic cells, but not Gatad2b, results in a deterministic naïve iPSC reprogramming where up to 100% of donor somatic cells successfully complete the process within 8 days. Genetic and biochemical analysis established a distinct sub-complex within the NuRD complex (Gatad2a-Chd4-Mbd3) as the functional and biochemical axis blocking reestablishment of murine naïve pluripotency. Disassembly of this axis by depletion of Gatad2a, results in resistance to conditions promoting exit of naïve pluripotency and delays differentiation. We further highlight context- and posttranslational dependent modifications of the NuRD complex affecting its interactions and assembly in different cell states. Collectively, our work unveils the distinct functionality, composition and interactions of Gatad2a-Chd4-Mbd3/NuRD subcomplex during the resolution and establishment of mouse naïve pluripotency.

## Introduction

Somatic cell reprogramming has boosted a major revolution in the field of stem cell research (Takahashi and Yamanaka, 2006). This is a technically simple process in which the induction of exogenous transcription factors, classically Oct4, Sox2, Klf4 and c-Myc proteins (abbreviated as OSKM), can induce somatic cells to convert back to embryonic pluripotent stem (ESC)-like cells, termed induced pluripotent stem cells (iPSCs) (Takahashi and Yamanaka, 2006; Wernig et al., 2007). Despite the relative simplicity of this process, conventional iPSC reprogramming is typically an inefficient and a-synchronized process, where less than 0.1-15% of the donor somatic cells undergo reprogramming over a period of 2-4 weeks (Hanna et al., 2009; Liu et al., 2016). Further, while donor somatic cells reprogram with different efficiencies, it is not possible to a priori definitively predict among individual identical donor somatic cells, which and when they will convert into iPSCs (Mikkelsen et al., 2008; Singhal et al., 2010). The latter attributes have supported the conclusion that conventional iPSC formation is a stochastic process, but amenable to acceleration by supplementing additional factors (Hanna et al., 2010; Ichida et al., 2014).

Multiple pioneering studies have devised alternative reprogramming protocols, some of which are donor cell specific, where rapid and up to 100% reprogramming efficiency can be obtained within a relatively short period (Bar-Nur et al., 2014; Lujan et al., 2015; Rais et al., 2013; Di Stefano et al., 2014; Vidal et al., 2014). Our group has found that controlled and partial reduction of Mbd3, a key component of Mbd3/NuRD (Nucleosome Remodeling and Deacetylation) corepressor complex, in concert with optimized OKSM delivery in naïve pluripotency conditions (2i/LIF, where 2i is applied 48-72 hours after reprogramming initiation, 5% O2 hypoxia conditions, and Vitamin-C-containing knock-out serum replacement), can lead to highly efficient and rapid iPSC formation with up to 100% reprogramming efficiency within 8 days (Rais et al., 2013). These high efficiencies and identity of iPSC cells generated have been independently validated using Cytof single cell analysis (Lujan et al., 2015), thus ruling out over-estimations from specific fluorescent pluripotency-associated reporters. In agreement with these findings, Grummt and colleagues showed that Mbd3 over expression blocks, whereas its depletion promotes, reprogramming of MEFs and partially reprogrammed cells (Luo et al., 2013a).

One of the major mechanisms by which Mbd3/NuRD repressive activity can be co-opted is through association with ectopically expressed transcription factors. The pluripotency factor Zfp281 directly recruits Mbd3/NuRD to repress Nanog promoter activity (Fidalgo et al., 2012). By directly interacting with many additional pluripotency promoting transcription factors in addition to Zfp281, including OSKM and Sall4, Mbd3/NuRD broadly suppresses reactivation of the pluripotency circuit (Fidalgo et al., 2011; Miller et al., 2016; Rais et al., 2013; Reynolds et al., 2012).

Essential to the promotion of reprogramming upon Mbd3 depletion is an optimal incomplete depletion of NuRD activity during a critical early reprogramming window (Rais et al., 2013). Both our group and Dos Santos et al. showed that using Mbd3^-/-^ somatic cells as starting material does not yield a boost in reprogramming (Rais et al., 2013; Dos Santos et al., 2014). In fact, as these cells can no longer proliferate *in vitro* after complete Mbd3 depletion, they trivially cannot reprogram as cell proliferation is indispensable for the iPSC process (Rais et al., 2013; Dos Santos et al., 2014). Further, the latter decrease in cell proliferation occurs prior and irrespective to OSKM induction, further supporting the notion that the inhibition of reprogramming efficiency simply results from hampering cell proliferation in the somatic state (Rais et al., 2013; Dos Santos et al., 2014), rather than establishing an epigenetic blockade for reprogramming per se. As such, Rais et al. focused mostly on using on Mbd3^flox/-^ cells and compared them to wild-type Mbd3^+/+^ cells. It should be noted that several iPSC boosting strategies rely on incomplete depletion of epigenetic modulators. For instance, Caf1, Sumo2, NCoR/SMRT and Ubc9 partial, but not complete, depletion was critical for maximal boosting of iPSC efficiencies by OSKM (Borkent et al., 2016; Cheloufi et al., 2015; Zhuang et al., 2018). Collectively, the latter observations further emphasize the need to better understand the NuRD components involved in obtaining the wanted effect of achieving deterministic iPSC reprogramming, which we set out to address in this study.

The NuRD co-repressive complex can assume multiple distinct complexes based on differential subunit composition in physiological and pathological states, including development, DNA damage response and cancer metastasis (Baubec et al., 2013; Le Guezennec et al., 2006; Spruijt et al., 2016; Yildirim et al., 2011). In addition to Mbd2 and Mbd3, which form two distinct mutually exclusive Mbd2/NuRD and Mbd3/NuRD complexes (Baubec et al., 2013; Le Guezennec et al., 2006; Xie et al., 2012), other canonical subunits include: Chd3 or Chd4 (chromodomain helicase DNA binding protein, also known as Mi2α and Mi2β, respectively), which retains an ATPase activity; RbAP48 and RbAP46, which are both histone-binding proteins; Hdac1 and Hdac2 (Histone deacetylase enzymes), which hold a deacetylation activity; Mta1, Mta2 and Mta3 (metastasis associated protein), which can interact directly with histones tails; Gatad2a or Gatad2b (also known as P66α and P66β, respectively) whose function remains to be fully defined (Alqarni et al., 2014). Quantitative mass spectrometry-based proteomics for Mbd3/NuRD complex has indicated that each NuRD unit contains six units of RbAP48 or RbAP46, three units of Mta1/2/3, two units of Gatad2a or Gatad2b, one unit of Chd3 or Chd4, and one unit of Hdac1 or Hdac2 (Smits et al., 2013). These results highlight the complexity and heterogeneity of NuRD complexes (Zhang et al., 2016). The latter is further complicated by the identification of context-dependent non-canonical components which include Doc-1, Zmynd8 (a zinc finger protein) and Lsd1 (Spruijt et al., 2010, 2016; Wang et al., 2009). Moreover, some of the proteins that contribute to the NuRD complex, maintain additional roles in the cell and take part in other complexes. For example, Chd4, which takes part in DNA damage response processes (Polo et al., 2010), is also part of an activating complex that includes CBP/p300 (Hosokawa et al., 2013). Likewise, Mbd3 has been reported to co-localize with aurora kinase at the mitotic spindle during mitosis and thus regulate cell cycle progression through undefined mechanisms that may possibly be NuRD independent (Sakai et al., 2002). These facts complicate assigning the outcome of perturbing components like Mbd3 or Chd4 to the NuRD complex exclusively, since observed functional changes might be stemming from perturbing only a certain sub complex of NuRD with a unique conformation or altering other different complexes that share some components with NuRD (e.g. Chd4 and Hdac1) (Sakai et al., 2002; Wang et al., 2009).

In this study, we set out to identify alternative strategies to block Mbd3 dependent NuRD activity while preserving somatic cell proliferation and viability. By dissecting the mechanisms and putative components of Mbd3/NuRD Subcomplex(es) relevant during early stages of reprogramming we have identified Gatad2a-Chd4-Mbd3 as a functional and biochemical axis underlying potent inhibition of re-establishment and maintenance of naïve pluripotency.

## Results

### Screening for alternative components to define and perturb Mbd3/NuRD sub complexes relevant for iPSC reprogramming

Optimized incomplete Mbd3 depletion during reprogramming has been shown to significantly improve boost iPSC formation efficiency (Luo et al., 2013a; Rais et al., 2013), but a complete reduction of this protein at the somatic stage results in cell cycle malfunctions (Rais et al., 2013) and, consequently, subsequent loss of reprogramming ability. In order to better understand the mechanisms of Mbd3 inhibitory effect on reprogramming, we adapted a transient silencing assay with siRNAs for screening other known NuRD components, aiming to identify those whose inhibition might dramatically enhance iPSC reprogramming while having minimal effect on somatic cell proliferation or viability, even if completely depleted at the protein level. To do so, we used a transgenic “secondary reprogramming” platform of MEF derived reprogrammable cell line, which already contains heterozygote TetO-inducible OKSM in the m*Col.1a* locus and heterozygote M2rtTA cassette in the *Rosa26* locus (Stadtfeld et al., 2010). We chose to examine canonical NuRD complex core members (Allen et al., 2013), with Mbd3 (used as a reference positive control), the mutually exclusive Mbd2, Chd4, Chd3, Mta2, Hdac2, Gatad2a and Gatad2b components **(Fig. 1A and S1A)**, and the non-canonical co-factor Zmynd8 **(Fig. S1D)**. siRNA was applied 48 hours and 96 hours after DOX administration (Reprogramming initiation). Consistent with our previous results, treating the cells with siRNA for Mbd3 and Chd4, but not Chd3 or Mbd2, has significantly improved the reprogramming rate as measured 9 days after DOX addition **(Fig. 1A-B, Fig. S1D)** (Cheloufi et al., 2015; Luo et al., 2013b; Rais et al., 2013). Remarkably, a significant and equivalent improvement in the reprogramming rate was seen in cells treated with siGatad2a (**Fig. 1A-B**). However, while depleting the NuRD components Mbd3 and Chd4 compromised the cell growth rate irrespective of inducing the Yamanaka OSKM factors (**Fig. 1C**), depleting Gatad2a had a relatively smaller effect on slowing proliferation rate of MEFs, as measured in cell growth assay (**Fig. 1C**) and BrdU incorporation (**Fig. 1D-E**). No significant increase in apoptosis in MEFs was evident either following depleting Gatad2a either (**Fig. 1F**).

**Figure 1.**
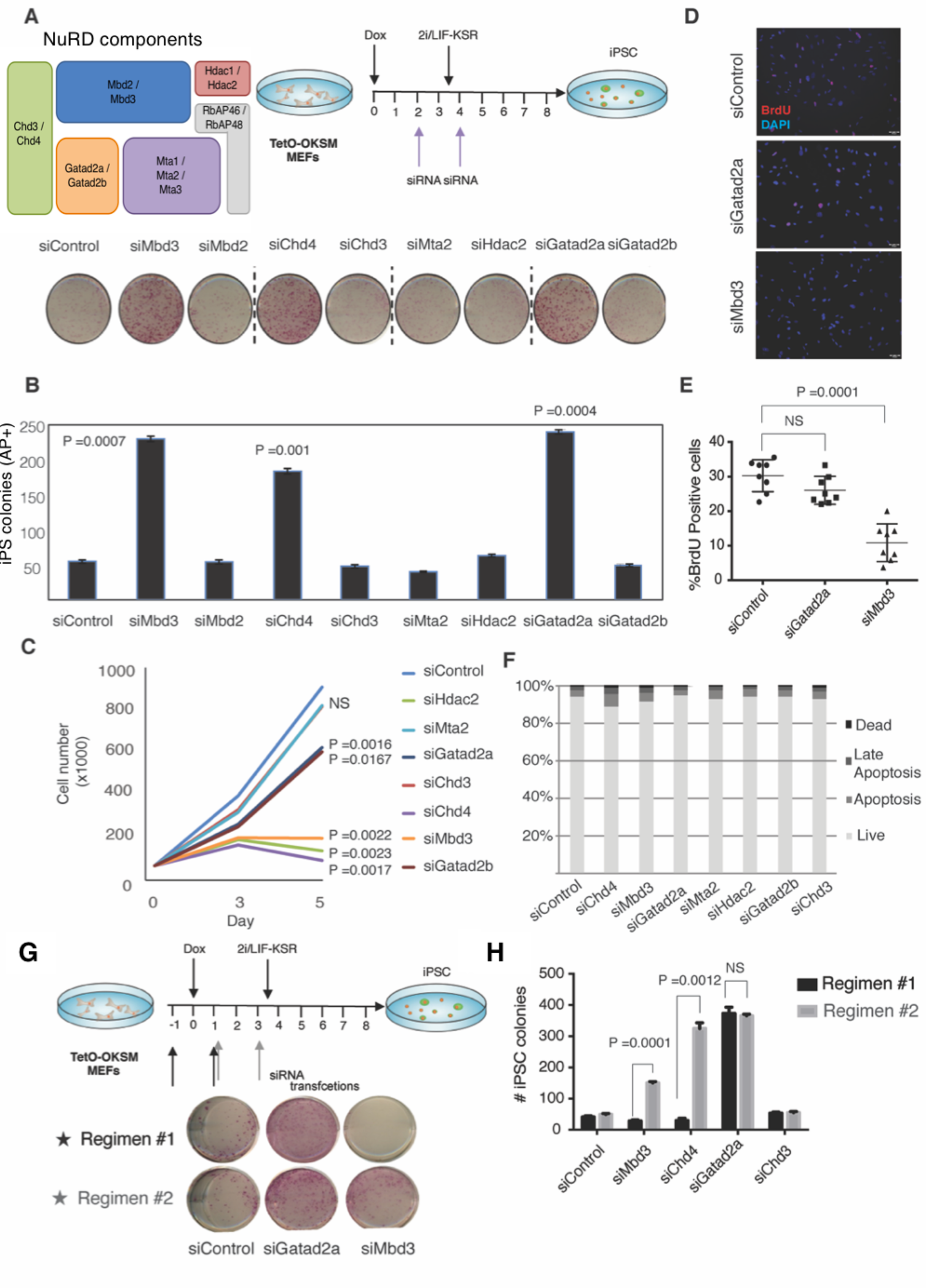
Gatad2a depletion increases reprogramming efficiency and does not ablate somatic cell proliferation. **A.** MEFs harboring TetO-OKSM and M2rtTA cassettes were transfected with siRNA targeting different canonical NuRD components (indicated in the illustration), 2 and 4 days after reprogramming initiation following DOX administration. Reprogramming was then evaluated by AP staining at day 8. **B.** Reprograming efficiency following siRNA treatments was evaluated using AP staining, after 8 days of reprogramming (n=3, two-sided Student’s t-test *p* values are indicated). **C.** Cell growth curves of MEFs treated with siRNA for the indicated NuRD components (two-sided Student’s t-test *p* values are indicated). Knockdown (KD) of Gatad2a, unlike KD of Chd4, Mbd3 and Hdac2, does not severely inhibit cell proliferation. **D.** Representative images of cells treated with siRNA targeting Mbd3 or Gatad2a and exposed to BrdU in order to evaluate proliferation. **E.** Quantitative evaluation of BrdU incorporation test, which shows normal proliferation in siScramble and siGatad2a, unlike in cells treated with siMbd3 (n=8, two-sided Student’s t-test *p* values are indicated Student’s t-test). **F**. Knockdown for different canonical NuRD components does not show a significant elevation in cell death or apoptosis. Viability and apoptosis induction were measured using FACS following Annexin-PI staining. **G.** Reprogramming efficiency following siRNA treatments targeting different NuRD components, at different time points. KD was performed at two distinct cycles: the early one (Regimen 1, marked in black) started one day prior to DOX induction, and the second one (Regimen 2, marked in grey) started one day post-DOX induction. **H.** iPSC reprogramming efficiency following different siRNA treatments. was evaluated at day 8. (n=3 per each condition, two-sided Student’s t-test *p* values are indicated). Gatad2a siRNA improves reprogramming whether administrated prior or post DOX administration, unlike siRNA targeting Mbd3 or Chd4, in which only in Regimen #2 iPSC colony formation efficiency was increased.

We next used modified regimens in order to evaluate the possibility of further optimizing the process, by repressing NuRD components at earlier time points during reprogramming. Chd4, Gatad2a, Chd3 and Mbd3 targeted siRNAs and siScramble were applied in two different transfection cycles: the first transfection cycle started in somatic cells before OKSM induction (pre-DOX), and the second one started 24 hours after reprogramming was initiated (post-DOX). In both regimens, a second transfection took place 48 hours after the first one in order to maintain optimal knockdown levels. siRNA mediated knockdown of Mbd3 and Chd4 achieved improved reprogramming only in the second cycle of transfection (post-DOX), while the siRNA transfection which started before OKSM induction (pre-DOX) had a negative effect on the reprogramming process, comparing to siScramble **(Fig. 1G-H)**. The latter is consistent with previous reports indicating that complete ablation of Mbd3 in the earlier somatic cell-like states hinders reprogramming due to higher sensitivity in blocking somatic cell proliferation (Rais et al., 2013; Dos Santos et al., 2014) which is essential for iPSC reprogramming (Hanna et al., 2009). However, as Gatad2a knockdown did not profoundly alter cell proliferation or viability, comparable enhancement in iPSC reprogramming rate was obtained in both transfection regimens **(Fig. 1G-H)**. iPSC lines acquired following siRNA treatment for Gatad2a were pluripotent as evident by uniform expression of pluripotency markers by immunostaining and generation of mature teratomas *in vivo* (**Supplementary Fig. S1B-C**). Taken together, these findings underscore Gatad2a as a potential alternative and technically more flexible way to inhibit Mbd3/NuRD repressive activity (i) early in the reprogramming process and (ii) without profoundly blocking somatic cell proliferation and viability when completely depleted.

### Gatad2a ablation facilitates deterministic iPSC reprogramming by OSKM

We next set out to more accurately examine the effect of abolishing Gatad2a during iPSC reprogramming by establishing multiple sets of genetically modified Gatad2a knock out (Gatad2aKO or Gatad2a^-/-^) secondary reprogrammable lines (Hanna et al., 2008; Hussein et al., 2014; Wernig et al., 2008) that were generated from defined parental isogenic Gatad2a^+/+^ wild-type (WT) cell lines. We first generated two different Gatad2a^+/+^ secondary reprogramming cell lines (**Fig. 2A and S2A)**: i) WT primary MEFs were reprogrammed with viruses encoding FUW-M2rtTA and STEMCCA-OKSM. A randomly picked iPSC clone was selected and rendered transgenic for a constitutively expressed mCherry allele (to allow tracking of viable somatic cells) and a stringent ΔPE-Oct4-GFP transgenic reporter for naïve pluripotency (Gafni et al., 2013; Rais et al., 2013). An isolated clone was then used as a WT control and as the source for an isogenically matched Gatad2a-/- null cell line. Notably, CRISPR/Cas9 was used to generate Gatad2a^-/-^ iPSCs/ESCs (**Fig. 2B-C**). Gatad2a^-/-^ clones were obtained with ˜40% successful targeting efficiency and were validated by PCR and Western blot analysis for ablation of Gatad2a (**Fig. S2B-D**). ii) Reprogrammable Gatad2a^+/+^ line was established by deriving an ES line from mice carrying Rosa26-M2rtTA^+/-^; m.Col1a-TetO-OKSM^+/-^; OG2-GOF18-ΔPE-Oct4-GFP allele (Boiani et al., 2004), and was then subjected to CRISPR/Cas9 Gatad2a KO strategy (**Fig. S2**). Isogenic WT and Gatad2a^-/-^ transgenic reprogrammable iPSCs/ESCS were injected into host blastocysts, and secondary MEFs were isolated from E12.5 embryos and were subjected to puromycin selection to eliminate nontransgenic host derived MEFs (**Fig. 2D and S3A**). In all experiments, isogenic sets were used side by side for comparative analysis (**Fig. 2A**). As previously optimized for Mbd3^flox/-^ donor cells (Rais et al., 2013), iPSC reprogramming was done in mES medium supplemented with DOX and 5% O_2_ conditions for the first three days, followed by changing to 20% O_2_ and knockout serum replacement (KSR) based medium supplemented medium with 2i/LIF (**Fig. 2A**). For all different lines tested, a profound improvement in the reprogramming rate and efficiency was observed, as the Gatad2a-KO cells achieve high rates alkaline-phosphatase positive colonies in 6 days only (**Fig. 2E)**. Reprogramming rates were also measured by FACS analysis by quantification of ΔPE-Oct4-GFP positive cells every 24 hours for 8 days without cell splitting or passaging to avoid any selective biases (**Fig. 2F, Fig. S3B**). Reminiscent of the kinetics measured in Mbd3^flox/-^ systems (Rais et al., 2013), ΔPE-Oct4-GFP starts to appear after 3 days of reprogramming, and its rates ascending prominently on day 4 and reach >95% after 8 days of OSKM induction (**Fig. 2E, Fig. S3B**).

**Figure 2.**
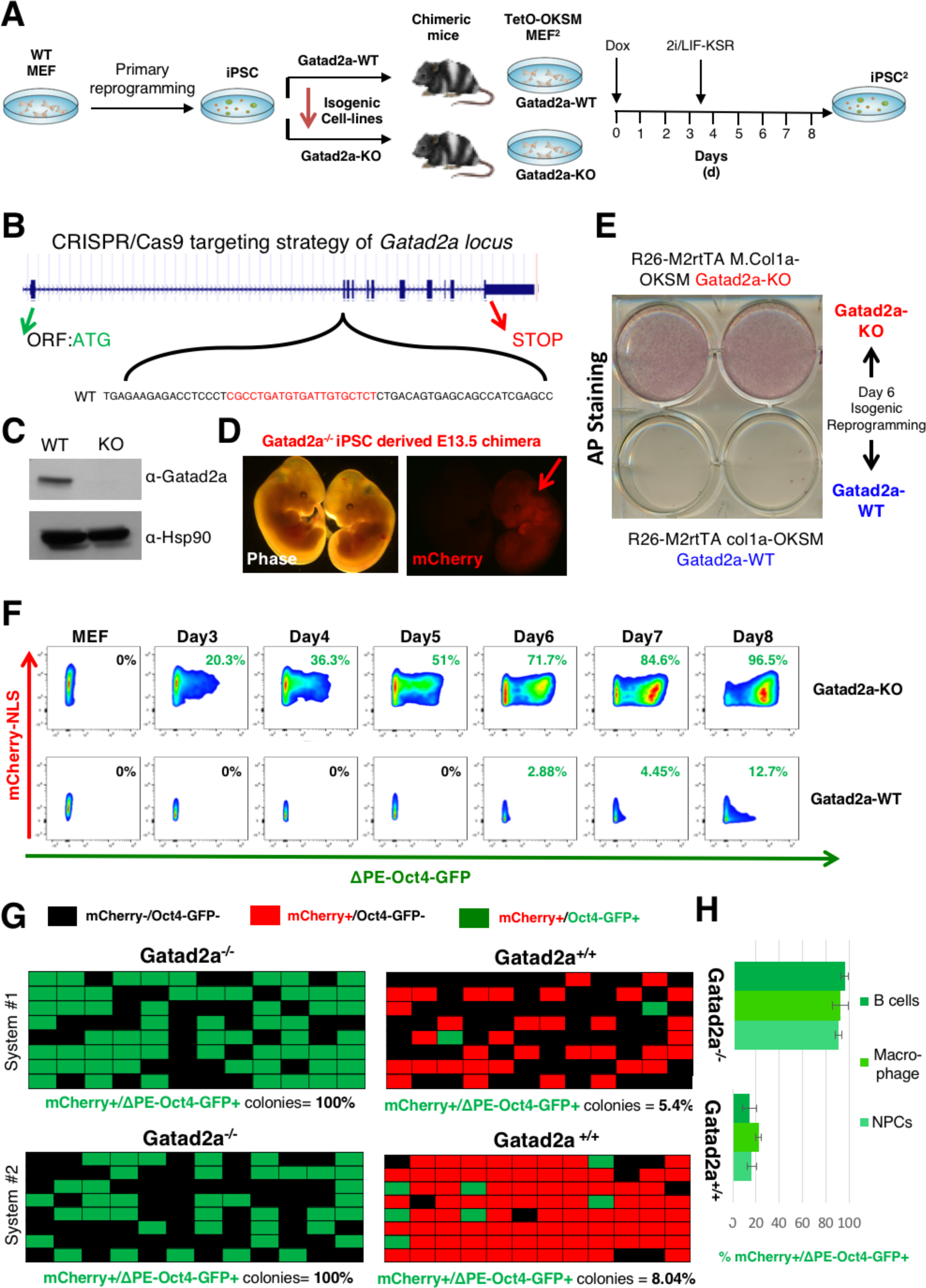
Deterministic induction of naïve iPSCs in Gatad2a-KO somatic cells. **A**. Scheme demonstrating strategy for generating secondary isogenic Gatad2a WT and KO lines, and comparing their reprogramming efficiency side by side. 2i/LIF-KSR conditions were introduced at day 3.5 during the 8-day course. More detailed information is provided in **Supplementary Fig. S2A** regarding different systems used herein. **B**. Targeting scheme of Gatad2a locus to generate knockouts by CRISPR/Cas9. **C**. Western blot validation of Gatad2a knockout in iPSC harboring M2rtTA and TetO-OKSM cassettes in comparison to its parental isogenic line. **D**. Representative images of Gatad2a^-/-^ iPSC derived E13.5 chimera. Red arrow highlights mCherry+ chimera which originates from mCherry labeled iPSCs that were microinjected. **E.** Bulk iPSC reprogramming as in **F**, but experiment was terminated after 6 days and iPSC colony formation was evaluated by Alkaline Phosphatase staining (AP+). **F**. Representative flow cytometry measurements of ΔPEOct4-GFP reactivation dynamics in polyclonal/bulk Gatad2a-WT and Gatad2a-KO isogenic cell lines. Throughout the course of the reprogramming experiment the cells were not passaged to avoid any biases. Reprogramming of secondary MEFs seeded as single cells. **G.** Representative summaries of single-cell iPSC reprogramming efficiency experiment. Secondary isogenic Gatad2a WT and KO reprogrammable MEFs carrying constitutively expressed mCherry-NLS and naïve pluripotency specific ΔPE-Oct4-GFP reporter were sorted and seeded as single-cell per well. Reprogramming was initiated by DOX administration according to panel **A**. Reprogramming efficiency was assessed after 8 days based on the number of wells in which mCherry+ cells formed an ΔPE-Oct4-GFP positive colony. Throughout the course of the reprogramming experiment the cells were not passaged to avoid any biases. **H.** The indicated secondary Gatad2a^+/+^ and Gatad2a^-/-^ somatic cell types were isolated and subjected to single cell reprogramming and evaluation of iPSC efficiency following 8 days of DOX. Reprogramming efficiency was assessed after 8 days based on the number of wells in which mCherry+ cells formed an ΔPE-Oct4-GFP positive colony. In summary, these results indicate that complete inhibition of Gatad2a (also known as P66a), a NuRD specific subunit, does not compromise somatic cell proliferation as previously seen upon complete Mbd3 protein elimination, and yet disrupts Mbd3/NuRD repressive activity on the pluripotent circuitry and yields 90-100% highly-efficient reprogramming within 8 days as similarly observed previously in Mbd3 hypomorphic Mbd3^flox/-^ donor somatic cells (Lujan et al., 2015; Rais et al., 2013).

We next utilized the fact that these lines harbor constitutive expression of nuclear mCherry marker that allows detection of viable somatic cells after single cell plating, and quantified reprogramming efficiency of MEFs at the single cell level. Gatad2a-KO MEF reproducibly yielded up to 100% iPS cell derivation efficiency by day 8 in two independent isogenic systems tested, as determined by ΔPE-Oct4-GFP expression in iPSC colonies obtained (**Fig. 2G)**. When isogenic Gatad2a WT cells reprogrammed under identical conditions, no more than 10% of clones reactivated ΔPE-Oct4–GFP in 8 days (**Fig. 2G**). In order to validate expression of pluripotency network, 96 well plates were fixed at day 8 and all wells were co-stained and found positive for endogenous Nanog/SSEA-1 and Oct4/Esrrb pluripotency marker expression in all clones tested at day 8, thus validating the authenticity of iPSC identity and reporters used herein (**Fig. S3C)**. Similar reprogramming efficiencies were observed upon reprogramming of other cell types including macrophages, neural progenitor cell (NPCs) and Pro-B cells (**Fig. 2H**).

We then applied microscopic live imaging of the reprogramming dynamics (Rais et al., 2013), by seeding somatic cells constitutively labeled with mCherry marker, and their reprogramming was evaluated by ΔPE-Oct4-GFP reactivation **(Fig. 3, Fig. S4 and Supplementary Video 1-2)**. Time-lapse measurements showed a dramatic increase in ES-like colony formation in Gatad2a^-/-^, 6 days after DOX induction: more than 98% of Gatad2a^-/-^ clonal populations reactivated ΔPE-Oct4-GFP pluripotency marker, compared to 15% in isogenic WT control cells, reprogrammed in identical growth conditions. Notably, while the most efficient conditions of reprogramming Gatad2a^-/-^ and Mbd3^flox/-^ cells involve applying 2i after 3 days (up to 100% at day 8), Gatad2a^-/-^ cells reprogram with dramatically higher efficiency at day 7 (40-70%) in all different conditions tested, compared to their isogenic WT controls **(Fig. S3D)**. Following Gatad2a depletion, we were not able to isolate stable partially reprogrammed cells that did not reactivate ΔPE -Oct4-GFP and could be stably expanded *in vitro* as typically can be obtained from OSKM transduced wild-type somatic cells (Borkent et al., 2016; Carey et al., 2010; Rais et al., 2013). The residual 1-15% ΔPE-Oct4-GFP negative colonies seen at day 6 in Gatad2a^-/-^ cells, rapidly become GFP+ within 1-3 days of continued reprogramming **(Fig. S3E)**.

**Figure 3.**
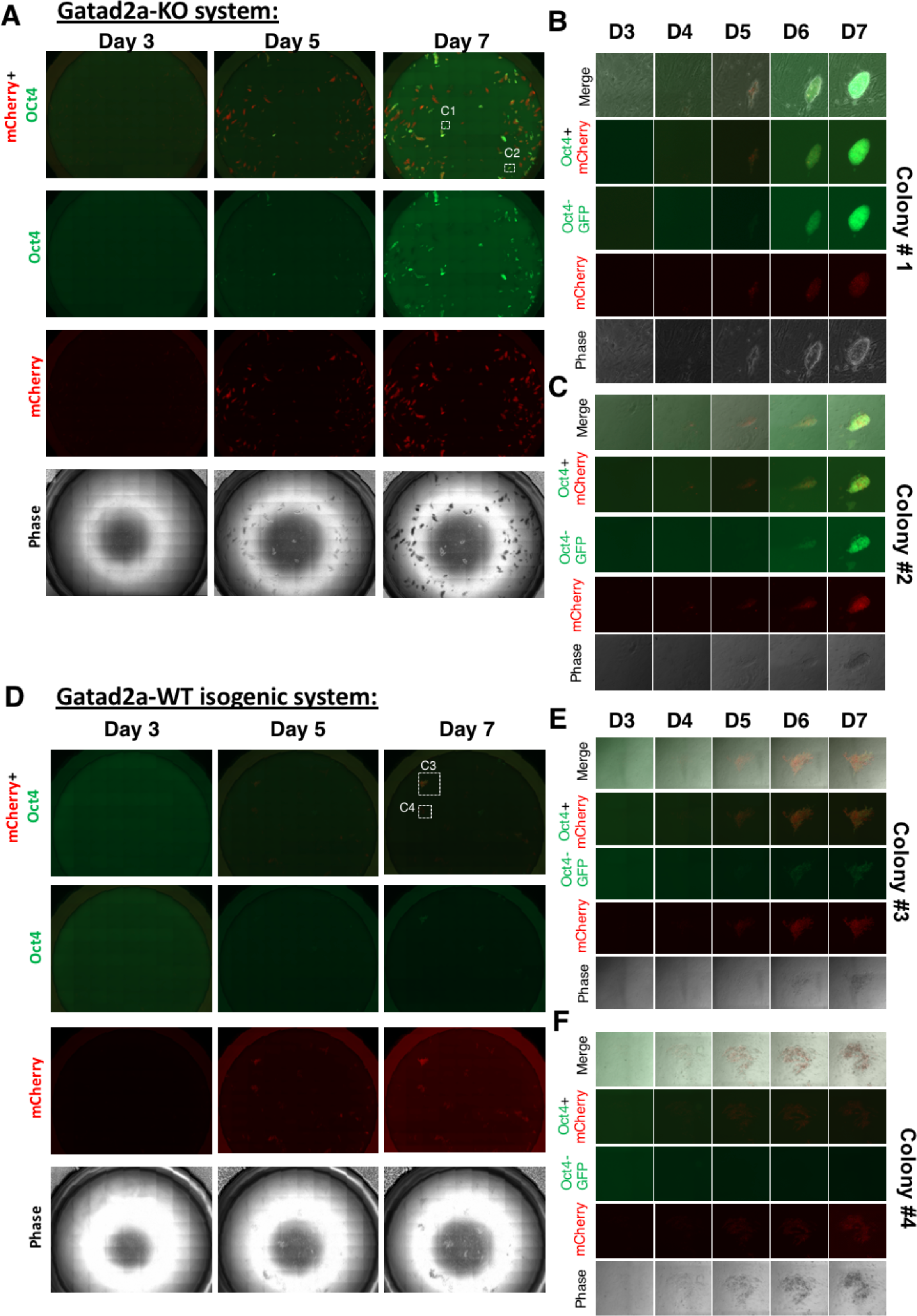
Whole-well mosaic live-cell imaging of reprograming following Gatad2a ablation. **A.** Selected time-points (Days 3, 5 and 7) of full-well mosaic, acquired during live imaging of Gatad2a-KO secondary MEF, harboring constitutive mCherry-NLS and ΔPE-Oct4-GFP reporter during OKSM mediated reprogramming. **B-C**. A zoom focusing on representative colonies (white squares C1 and C2 respectively in panel **A**), showing multiple time-points from live-imaging of the colonies’ reprogramming. The global and focused images show how nearly all donor cells become ΔPE-OCT4-GFP+. **D.** Selected time-points of full-well mosaic, acquired during live-imaging of Gatad2a-WT (isogenic control to cells used in **A-C**) reprogramming. **E-F**. A zoom focusing on representative colonies (white squares C3 and C4 respectively in panel **D**), showing multiple time-points from live-imaging of the colonies’ reprogramming. The global and focused images show how most growing clones (labeled with mCherry) do not reactivate ΔPE -Oct4-GFP expression (e.g. clone #3 in panel E), and only some rare clones become iPSCs (e.g. Colony #4 highlighted in panel **F**). Please see **Supplementary Videos 1-2**, from which these images were taken.

Twelve Gatad2a^-/-^ derived iPSC lines from each of the two described secondary reprogramming systems were established independently of DOX and passaged before subsequent functional characterization. All randomly tested clones stained positive for alkaline phosphatase (AP), Oct4 and Nanog pluripotency markers **(Fig. S3F)**. Mature teratomas and high contribution chimeras were obtained from multiple iPSC clones consistent with adequate reprogramming to ground state pluripotency **(Fig. S5A-B)**. To evaluate the molecular extent and authenticity of reprogramming in OSKM transduced Gatad2a^+/+^ and Gatad2a^-/-^ cell, we conducted global gene expression analysis on bulk donor MEFs at days 1-8 following DOX induction without cell passaging or sorting based on pluripotency reporter expression and compared them to iPSC and ESC lines. For all mouse iPSC reprogramming experiments dedicated for genomic analysis, irradiated human foreskin fibroblasts were used as feeder cells, as any sequencing input originating from the use of human feeder cells cannot be aligned to the mouse genome and is therefore omitted from the analysis. Gatad2a^-/-^ somatic cells and not Gatad2a^+/+^ cells clustered separately from donor fibroblasts already at day 1 following DOX **(Fig. 4A-B)**. Importantly, Gatad2a^-/-^ MEFs cluster together with WT MEFs, thus ruling out a possibility of dramatically different transcriptional state before DOX **(Fig.4A-B, Fig. S5C, Fig S2E)**. Remarkably, by day 8 Gatad2a^-/-^ cells were transcriptionally indistinguishable from multiple ESC and sub cloned established iPSC lines **(Fig. 4A-B).** Key regulators of mouse fibroblasts or mouse pluripotency were similarly expressed in Gatad2a^-/-^ and Gatad2a^+/+^ cell lines **(Fig. 4A-B, Fig. S5C)**. Genome wide chromatin mapping for H3K27acetyl (K27ac) by Chromatin Immunoprecipitation followed by sequencing analysis (ChIP-seq), also confirmed that by day 8, only Gatad2a^-/-^ transduced MEFs had assumed an ESC-like chromatin profile **(Fig. 4C)**. Genome wide DNA methylation mapping by whole genome bisulfite sequencing (WGBS) confirmed that the Gatad2a^-/-^ polyclonal population of MEFs and iPSCs are positively correlated to their WT counterparts **(Fig. 4C)**. More extensive comparison to Mbd3^flox/-^ system, shows they are also comparable in their RNA-Seq, H3K27ac and WGBS to Mbd3^flox/-^ counterpart samples (Zviran et al., 2017)(Accompanying related manuscript with PDF co-submitted). Moreover, FISH experiments for asynchronous DNA replication show that cells undergoing reprogramming start from asynchronous replication (in their somatic state) but adopt synchronous replication pattern which suits ground state naïve pluripotent stem cells (Masika et al., 2017) as early as day 5 (**Fig. S5H**), coinciding with the robust reactivation of ΔPE-Oct4-GFP reporter in the massive majority of donor cells. Collectively, the above results indicate that Gatad2a depletion following OSKM induction in optimized 2i/LIF containing conditions yields authentic molecular reestablishment of the ground state of pluripotency in the entire population of donor somatic cells and their progeny.

**Figure 4.**
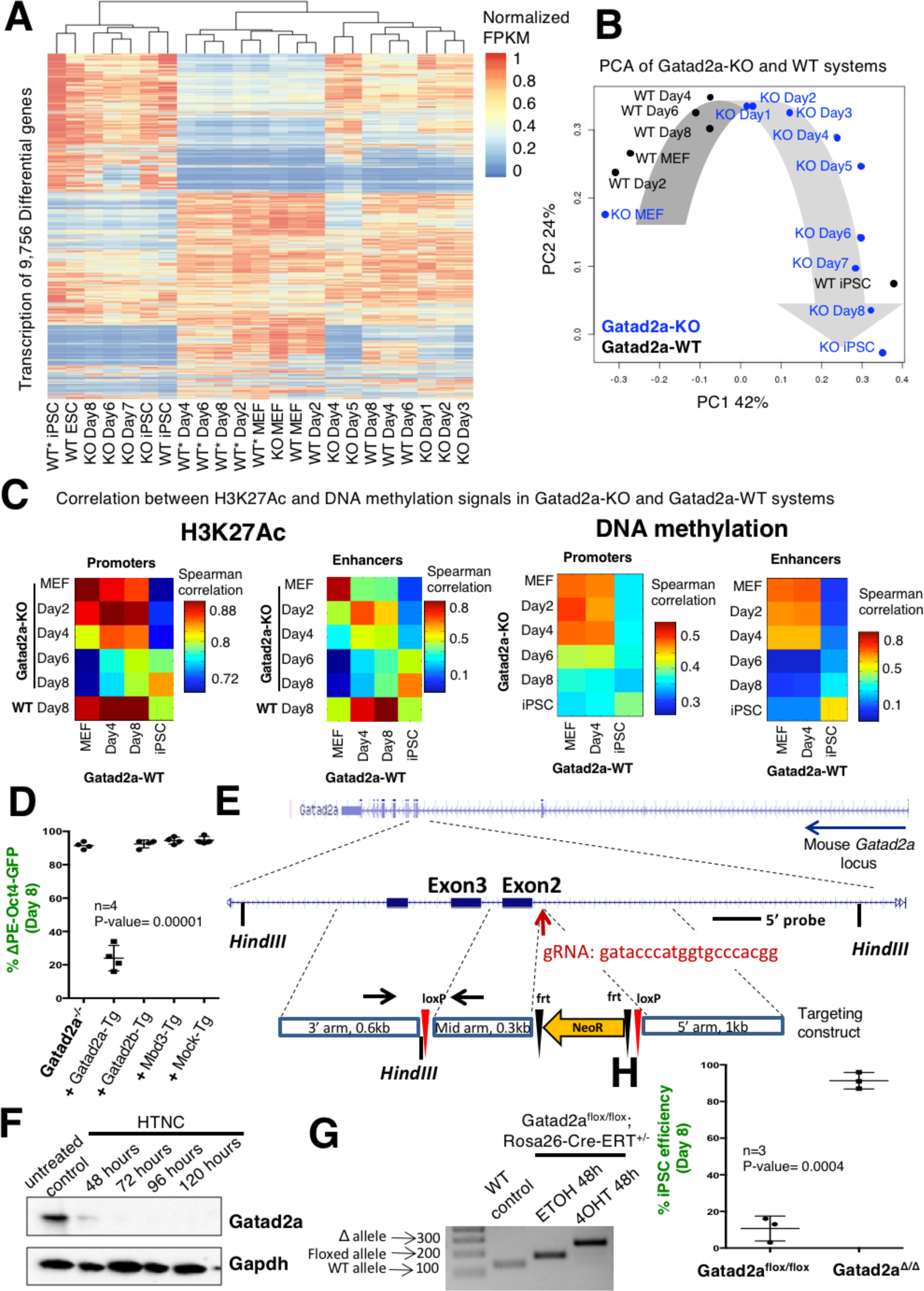
Authentic naïve pluripotency establishment by OSKM specifically following Gatad2a depletion. **A.** Hierarchical clustering of transcription profiles of samples from Gatad2a^-/-^ system and two WT systems, where WT are isogenic to Gatad2a-/- and WT* are non-isogenic. Values are unit-normalized FPKM (see **Methods**). IPSC samples from all systems cluster together, along with Gatad2a^-/-^ samples from days 6-8. MEF samples from all system also cluster together, along with all WT samples. Starting Gatad2a^-/-^ MEF and IPS cells are comparable to those of WT. **B.** PCA of Gatad2a^-/-^ and isogenic Gatad2a^+/+^ WT system, showing the transformation from MEF to iPSC, and that MEF and iPSC are similar in both systems. PCA was calculated over the same set of genes shown in (**A**). Only isogenic sample sets were included in analysis shown in Panel **B**. **C**. Correlation matrices between Gatad2a^-/-^ and WT cells, based on H3K27ac signal (left), and DNAmethylation signal (right). Correlation of H3K27ac signal was calculated over promoters of all differential genes (n=14,033), or over differential enhancers (n=12,153). Correlation pf DNA methylation signal was calculated over promoters of differential genes that are covered by the sequencing method (n=9,134), or over covered differential enhancers (n=11,413). **D**. Reprogramming efficiency of Gatad2a^-/-^ secondary cells was evaluated before and after introducing different exogenous transgenes (Tg), including Gatad2a-Tg**. E.** Targeting strategy for generating Gatad2a conditional knockout reprogrammable system and ESCs. **F.** Western-blot time-course based validation for Gatad2a protein depletion in Gatad2a^flox/flox^ MEFs following Tat-Cre treatment (HTNC). **G.** Complementing PCR based validation of Gatad2a floxed allele deletion following 4OHT treatment (the cells also harbor a correctly targeted Rosa26-CreERT allele). **H.** Gatad2a^fl/fl^ reprogrammable MEF were derived and underwent Cre induction for 4 days, and only afterwards DOX reprogramming was initiated for 8 days as described in **Fig. 2A, E** (n=3 per conditions, student’s t-test *p* value is indicated).

Of note, inhibiting Gatad2a expression was not sufficient to induce iPSC formation in the absence of exogenous OSKM overexpression in somatic cells (**Fig. S5D**). Gatad2a ablation did not replace exogenous OSK expression neither to give rise to iPSCs and the presence of exogenous cMyc was essential to obtain iPSCs at up to 100% efficiency following factor transduction within 9 days (**Fig. S5D**). Last, in order to validate the specificity of Gatad2a function, we reconstituted Gatad2a expression in Gatad2a^-/-^ MEFs using viral infection of Gatad2a under FUW-TetO promoter. Subsequent reprogramming resulted in a significant decrease of reprogramming efficiency comparing to Gatad2a^-/-^ cells (**Fig. 4D and Fig. S5E-G)**. Notably, Gatad2b over expression failed to achieve the same phenotype and had no influence on reprogramming efficiency of Gatad2a^-/-^ MEFs. These results suggest a non-redundant role for Gatad2a which cannot be compensated for by Gatad2b (**Fig. 4D and Fig. S5E-G)**. The latter is consistent with recent observation that Gatad2a and Gatad2b form mutually exclusive complexes (Spruijt et al., 2016) and the fact that Gatad2a^-/-^ mice are embryonically lethal by E8.5-E10.5 despite of intact residual Gatad2b expression (Marino and Nusse, 2007).

Finally, while Gatad2a^-/-^ MEFs had epigenetic and functional features of somatic WT MEFs (including RNA-seq, WGBS and H3K27ac) (**Fig. 4A-C, Fig S2E**), in order to fully exclude a possibility that a boost in iPSCs efficiency is partially reliant on incomplete differentiation of somatic cells generated while Gatad2a was completely absent during the differentiation process, we generated Gatad2a^flox/flox^ Secondary iPSCs (**Fig. 4E**). Subsequently, the safe harbor *Rosa26* locus was correctly targeted with a knock-in CRE-Ert allele. These cells were microinjected into host chimeras and secondary Gatad2a^flox/flox^ MEFs were derived and used for reprogramming efficiency measurement with the presence or absence of Cre recombinase or Tamoxifen (4OHT) which were both efficient in ablating Gatad2a protein expression (**Fig. 4F-G**). Reprogramming efficiency and kinetics from Tat-CRE (HTNC) treated cells (Gatad2a^Δ/Δ^) were indistinguishable from Gatad2a^-/-^ cells, while reprogramming efficiency of isogenic paternal isogenic Gatad2a^flox/flox^ somatic cells were below 18% (**Fig. 4H**).

### Gatad2a restrains naive pluripotency maintenance and lineage priming of mouse ESCs

Mbd3 has been shown to inhibit naïve pluripotency expression, and in its absence, mouse ESCs become more stable under naive conditions and exhibit significant kinetic delay to undergo differentiation (Kaji et al., 2006, 2007; Reynolds et al., 2012). In order to evaluate Gatad2a influence on murine naïve pluripotency maintenance, Gatad2a was knocked out in a murine ESC line harboring the OG2 ΔPE-Oct4-GOF18-GFP transgenic reporter that specifically marks the naïve pluripotent state (Bao et al., 2009; Yoshimizu et al., 1999) (**Fig. 5A**). The Gatad2a^-/-^ ESC stained positive for all pluripotency markers tested (**Fig. 5B)** and exhibited a normal protein expression profile **(Fig. 5C)**, suggesting that Gatad2a is dispensable for naïve pluripotency maintenance. ΔPE-Oct4-GFP reporter levels were similar in WT and Gatad2a-KO, as shown by FACS analysis and representative pictures of colonies from both cell lines (**Fig. 5A-B, D)**. Gatad2a-/-, but not WT cells, could be maintained in serum only (FBS only) conditions without supplementation of LIF for >12 passages without any compromise in ΔPE-Oct4-GFP expression levels (**Fig. 5D**), consistent with an increase in their naïve pluripotency stability and similar to previously obtained results with Mbd3^-/-^ murine ESCs (Kaji et al., 2007; O’Shaughnessy-Kirwan et al., 2015; Rais et al., 2013).

**Figure 5.**
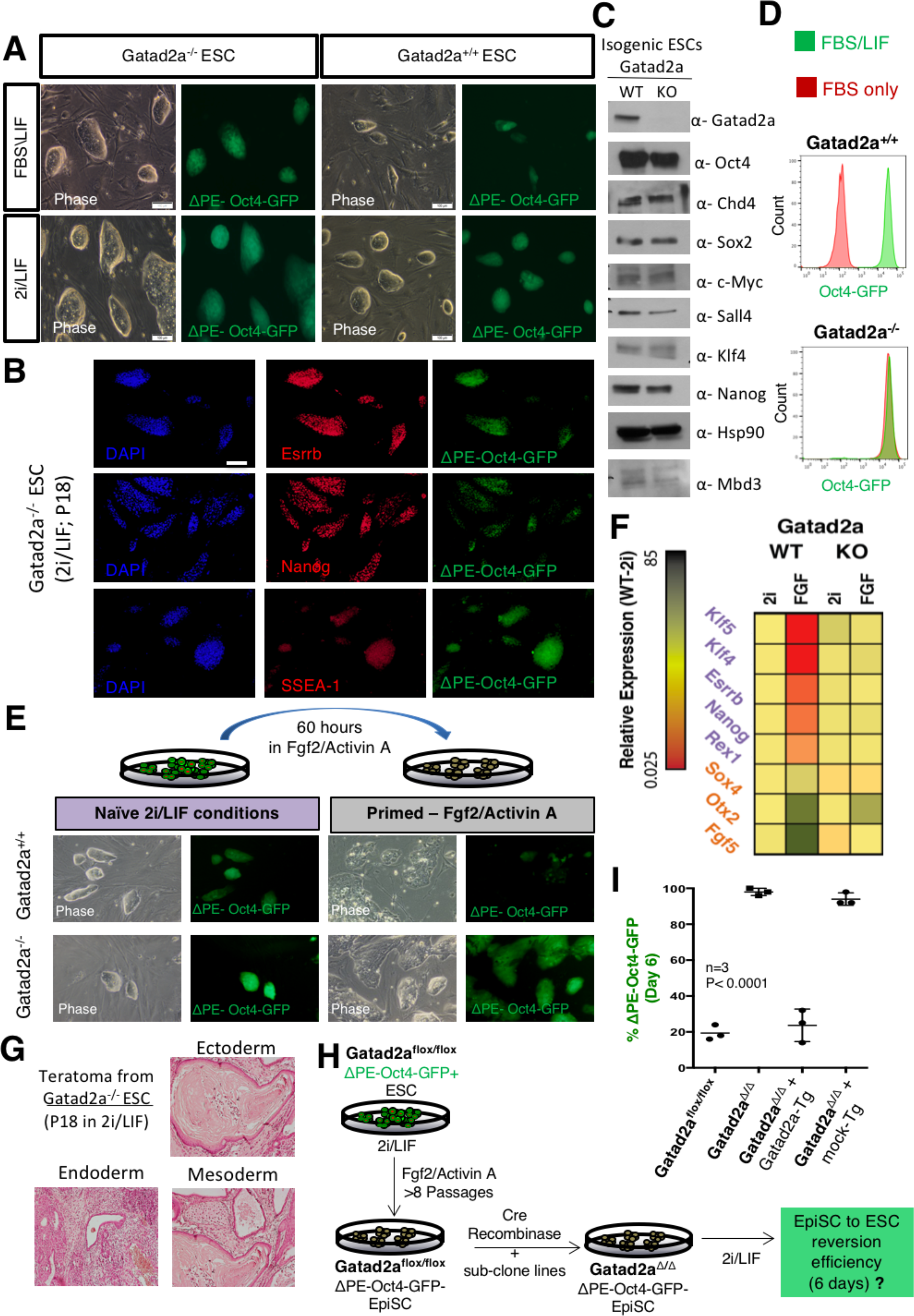
Gatad2a depletion enhances naïve pluripotency maintenance and induction from primed EpiSCs. **A.** Representative images of isogenic Gatad2a^+/+^; #PE-Oct4-GFP and Gatad2a^-/-^ ΔPE-Oct4-GFP ESCs that can be stably expanded in both FBS/LIF and 2i/LIF naïve conditions. (scale= 100 μM). **B**. Representative immunostaining of Gatad2a-KO ES. The Cells stained positive for various pluripotency markers, including Esrrb, Nanog and SSEA-1. (Scale bar = 100 μM). **C**. Western blot comparing pluripotency proteins’ level between Gatad2a-WT and KO isogenic ES lines. **D.** Isogenic Gatad2a^+/+^ and Gatad2a^-/-^ ESCs were maintained on Gelatin coated plates in FBS/LIS or FBS only conditions for 5 passages and then subjected to FACS analysis for OG2 ΔPE-Oct4-GFP pluripotency levels. **E**. Priming of Gatad2a^+/+^ and Gatad2a^-/-^ naïve ESCs harboring OG2 ΔPE- -Oct4-GFP reporter by changing conditions from 2i/LIF to Fgf2/Activin A was performed. While WT cells exhibit dramatic morphological change after 60 hours of treated with primed Fgf2/Activin A containing medium, Gatad2a-KO show resistance in losing their domed morphology and downregulating naïve pluripotency specific ΔPE-Oct4-GFP reporter. **F.** Transcriptional expression of different pluripotency and early differentiation markers before and after Fgf2/Activin A exposure, in isogenic Gatad2a^+/+^ and Gatad2a^-/-^ ESCs, presented as a relative expression column scheme. **G.** Gatad2a-KO ESCs generate mature and normal teratomas, including mature cells from all three germ layers of development. **H.** Strategy for generating isogenic Gatad2a^flox/flox^ ΔPE-Oct4-GFP was established from parental cells expanded for 8 passages in FGF2/Activin A and was validated for priming by FACS and RT-PCR analysis. Established Gatad2a^flox/flox^; ΔPE-Oct4-GFP was treated with Tat-CRE, and sub cloned isogenic Gatad2a^Δ^/# EpiSC lines were derived and used for analysis within additional 5 passages of their sub cloning. **I**. Single cell reprogramming efficiency and quantification for EpiSC reprogramming from different mutant EpiSC lines. Rescue Gatad2a-Tg or control Mock-Tg were ectopically expressed in Gatad2a^Δ^/Δ EpiSC and included in the analysis. (n=3 per time point, *p* Value<0.0001, two-sided Student’s t-test).

We next set out to find whether Gatad2a deletion has a functional effect on the cell ability to undergo lineage priming and differentiation, by examining conversion of naïve ESC into primed Epiblast like cells (EpiLC) *in vitro* (Hayashi et al., 2011) after 60 hours expansion in primed Fgf2/Activin A defined conditions (**Fig. 5E**). Gatad2a^+/+^ cells showed a rapid decrease in ΔPEOct4-GFP signal and lost their domed-like shape morphology, while Gatad2a^-/-^ cells retained their naïve morphology and had a negligible decrease in ΔPE-Oct4-GFP following 60h of priming in bFGF/Activin A conditions **(Fig. 5E)**. Consistently, RT-PCR analysis revealed that conversion of Gatad2a^-/-^ caused a smaller downregulation of naïve pluripotency related genes such as Esrrb, Nanog, Rex1 and Klf4, compared to isogenic wild-type control cells (**Fig. 5F, Fig. S6A)**. In addition, KO cells were deficient in up-regulating early development genes such as Otx2 and Fgf5 (**Fig. 5F, Fig. S6A**). Notably, this delay in undergoing priming is not infinite and is resolved later on, as Gatad2a^-/-^ are able to differentiate despite of the latter delay and form mature teratomas upon microinjection into immune deficient mice *in vivo* **(Fig. 5G)** as similarly described for Chd4^-/-^, Mbd3^-/-^ and hypomorphic Mbd3^flox/-^ ESCs. Collectively, these observed phenotypes resemble those previously validated with Mbd3 and Chd4 depleted naïve ESCs (Kaji et al., 2007; O’Shaughnessy-Kirwan et al., 2015; Rais et al., 2013), and once again underlines the common functionality of these three NuRD components.

To evaluate whether Gatad2a depletion promotes reversion of primed cells to naïve pluripotency, Epiblast stem cells (EpiSCs) (Brons et al., 2007; Greber et al., 2010; Najm et al., 2011)were established and validated from Gatad2a^flox/flox^ ESCs that harbor naïve specific OG2 ΔPEOct4-GFP reporter (**Fig. 5H**). In comparison to isogenic Gatad2a^flox/flox^ EpiSCs, single cell clonal analysis for epigenetic reversion of EpiSCs demonstrated >95% ΔPE-Oct4-GFP+ single cell reversion efficiency in Gatad2a depleted cells **(Fig. 5I)**. These efficiencies are similar to those obtained following reversion of Mbd3^flox/-^ EpiSCs (Rais et al., 2013). Further, deleting Gatad2a in EpiSCs while maintaining them in primed Fgf2/Activin A conditions for extended passages (>10) (**Fig. S6B**) partially compromised their primed identity towards naivety. While ΔPE-Oct4-GFP remained negative in both early and late passage Gatad2a^Δ/Δ^ cells expanded in primed conditions, late passage Gatad2a^Δ/Δ^ had reduced expression of primed markers such as Brachyury (T) and upregulated expression of Nanog and Esrrb naïve markers in comparison to early passage Gatad2a^Δ/Δ^ cells (P3-5) (**Fig. S6C-D)**. These results directly demonstrate that reduction of Gatad2a protein levels compromises the identity of the primed pluripotent state and renders nearly complete reversion of all donor primed EpiSCs to ground state pluripotency upon exposure to robust naïve pluripotency conditions.

### Mbd3, Gatad2a and Chd4 form a key molecular axis within the NuRD complex that restrains naïve pluripotency

Gatad2a inhibition was a key and dominant contributor to the radically efficient regression towards naïve pluripotency reported herein (**Fig. 2**). Thus, we aimed to define the mechanisms by which Gatad2a influences Mbd3/NuRD to facilitate inhibition of naïve iPSC reprogramming. In order to do this, we established an ESC line that harbors a DOX inducible allele of 2XFlag-Mbd3 (Rosa26-M2RtTA^+/-^ Col1a:TetO-2XFlag-Mbd3^+/-^) **(Fig. S2Aiii)** (Hochedlinger et al., 2005). Subsequently, we generated an isogenic Gatad2a^-/-^ clone by CRISPR/Cas9 mediated targeting (**Fig. S2**). In order to estimate the influence of Gatad2a-KO on Mbd3 interactome we applied LC-MS/MS analysis of Mbd3 protein interactions in Gatad2a-KO line and its isogenic WT control. When focusing the analysis on NuRD components, we noted that all canonical members of the NuRD complex were identified in both samples in high abundance and high correlation, except for Gatad2a and Chd4, which were missing from the Gatad2a-KO sample (**Fig. 6A**).

**Figure 6.**
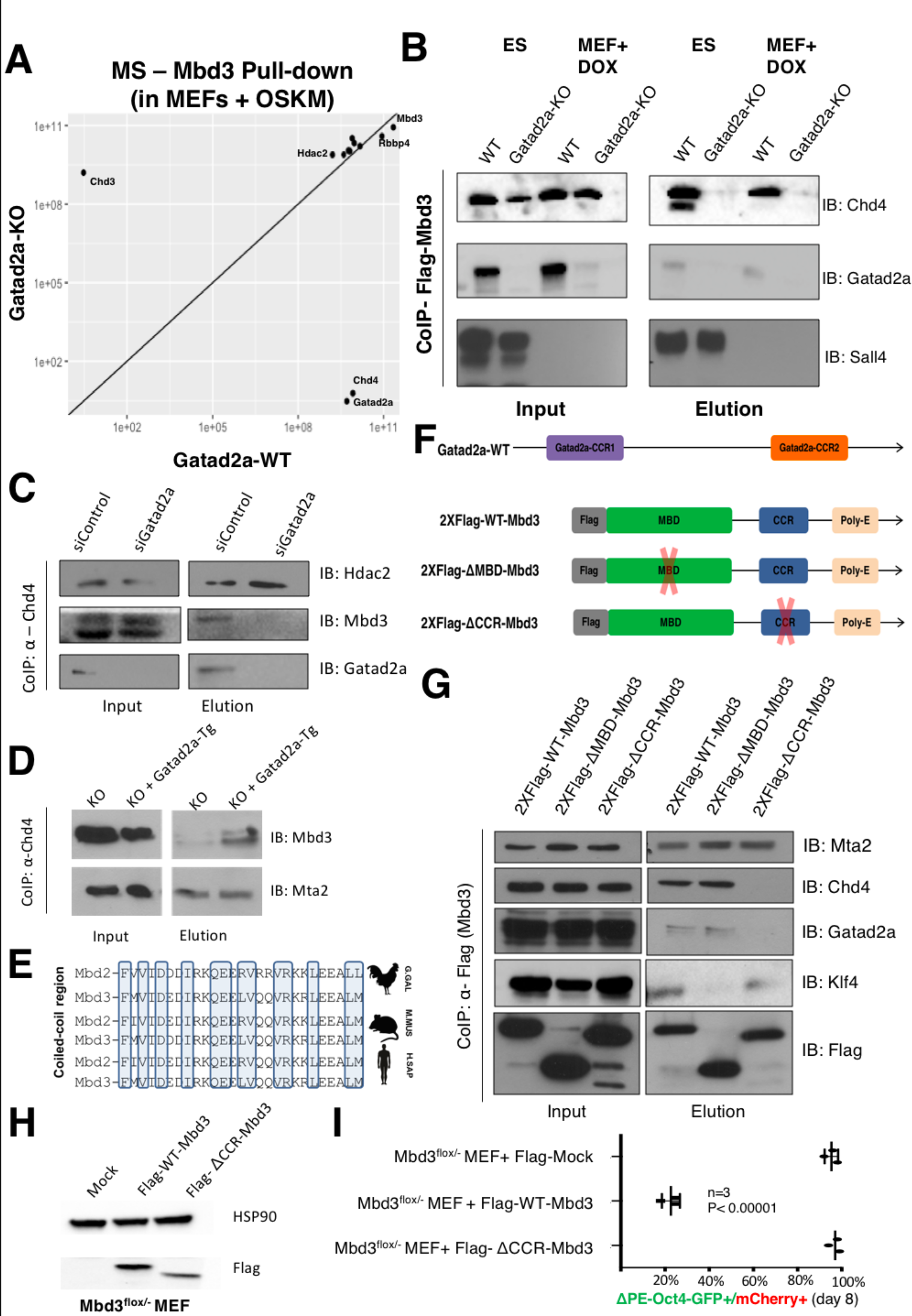
Gatad2a, Mbd3 and Chd4 constitute a critical axis within the NuRD complex mediating iPSC reprogramming inhibition. **A.** Mass spectrometry analysis of NuRD complex components binding efficiency to Mbd3 in Gatad2a-KO and its isogenic Gatad2a-WT line, in MEF cells during reprogramming by OSKM. Only putative NuRD components are presented and can be seen in an equal strength at both platforms, except for Chd4, which does not bind Mbd3 in Gatad2aKO. Flag-Tagged Mbd3 was used to establish a platform for studying Mbd3-binding proteins, by correct targeting of TetO-Mbd3-Flag into the M. Col1a locus. Isogenic Gatad2a-KO were generated from this line with CRISPR/Cas9 and used for the IP and MS analysis indicated above (See **Fig. S2Aiii**). **B.** Flag-Mbd3 CoIP in Gatad2a-WT and Gatad2a-KO cells. Experiments were conducted both in MEF cells and ESs. **C.** Cells during reprogramming were treated with siRNA targeting Gatad2a, and pellets collected after four days. CoIP of Chd4 shows that siGatad2a prevents Mbd3 binding to Gatad2a but also to Chd4. **D.** Gatad2a-KO MEF with overexpression of Gatad2a (transgenic recovery; abbreviated as Gatad2a-Tg) or Mock were subjected to Chd4-CoIP. Gatad2aoverexpression recovers the binding of Chd4 to Mbd3 and does not affect its binding to Mta2. **E.** The coiled coil region of Mbd3 is highly conserved between different organisms, and different proteins in the MBD family. The highlighted amino acids are crucial for Gatad2a binding to Mbd2 or Mbd3. **F.** A scheme of Flag tagged Mbd3, and mutant forms of Mbd3: lacking the Coiled coil region (#CCR-Mbd3) or the methyl-binding domain (#MBD-Mbd3). **G.** WT-Mbd3 and both mutants were over-expressed in 293T cells and were subjected to Flag-Mbd3 CoIP to examine their protein interactions. While the deletion of MBD prevents the binding of Klf4, only the deletion of the coiled coil region abolished the binding to Chd4 and Gatad2a. **H-I.** Mbd3^fl/-^ secondary MEF, harboring #PE-Oct4-GFP reporter, were transfected with two different forms of Flag tagged WT-Mbd3 and ΔCCR-Mbd3. The cells were then subjected to reprogramming. While WT-Mbd3 significantly reduced reprogramming efficiency (*p* Value<0.0001, two-sided Student’s t-test, n=3), ΔCCR-Mbd3 expression was not able to inhibit deterministic reprogramming in the cells, consistent with its inability to interact and recruit Chd4 to the assembled complex.

In light of the above, we set out to establish whether and how Mbd3, Chd4 and Gatad2a constitute a biochemical axis within the NuRD complex. Indeed, co-immunoprecipitation (Co-IP) analysis for Mbd3 in ESCs and MEFs showed that in the absence of Gatad2a, Chd4 could not be immunoprecipitated with Mbd3 (**Fig. 6B**). The latter principle was validated following IP of endogenous Chd4 in MEFs expressing OSKM undergoing reprogramming, showing that upon depletion of Gatad2a via siRNA, Chd4 can no longer directly interact with Mbd3 (**Fig. 6C**). Transgenic ectopic reconstitution of Gatad2a in Gatad2a^-/-^ MEFs (KO + Gatad2a-Tg), reestablished specific interaction between Mbd3 and Chd4 (**Fig. 6D**).

We next wanted to determine which domains of Gatad2a and Mbd3 regulate this specific interaction. Mbd3, which is a member of the Methyl-binding-domain (MBD) proteins retains two domains – the Methyl binding domain, which has been shown to mediate its interaction with transcription factors like OSKM during iPSC reprogramming (Rais et al., 2013), and the highly conserved Coiled coil region whose precise function in Mbd3 is not fully characterized (**Fig. 6E**). 2xFlag-tagged Mbd3 (2XFlag-WT-Mbd3) expression vector was generated together with two mutant versions of Mbd3, one of which lacks the coiled-coil region (2xFlag-ΔCCR-Mbd3) and the other one lacks the Methyl binding domain (2XFlag-ΔMBD-Mbd3) **(Fig. 6F).** Co-IP experiments showed that while the elimination of the coiled-coil region of Mbd3 (2XFlag-ΔCCR-Mbd3) does not interrupt binding to reprogramming factors (e.g. Klf4), it does prevent exclusively Gatad2a and Chd4 binding **(Fig. 6G)**. On the contrary, ΔMBD-Mbd3 was unable to bind Klf4, but maintained its ability to bind to different NuRD components, like Gatad2a and Chd4, as similarly shown in previous publications (Rais et al., 2013) **(Fig. 6G).** Reconstitution of Mbd3^flox/-^ cells with ΔCCR-Mbd3 transgene, which is missing the coiled-coil domain, did not inhibit reprogramming efficiency, consistent with its inability to recruit Chd4 (**Fig. 6H-I**).

In order to further characterize Mbd3-Gatad2a-Chd4 axis, we tried to interrupt the axis assembly by creating a “competition” over the binding site of Gatad2a to Mbd3. We over-expressed Gatad2a-coiled coil region (Gatad2a-CCR1) in a truncated construct (**Fig. S7A-B**) or as a short peptide (**Fig. S7A, C-D**) together with 2XFlag-Mbd3. Indeed, the over-expression of Gatad2a Coiled-coil region reduced the endogenous Gatad2a and Chd4 binding to Mbd3 (**Fig. S7B-D**). Finally, overexpression of the MBD domain of Mbd3 as an independent peptide directly interacted with Oct4 and Hdac2, showing again that the MBD is required and sufficient for the binding of the pluripotency factors (Rais et al., 2013), but not to Gatad2a and Chd4 (**Fig. S7E**). Collectively these results establish a function of Mbd3-CCR as the mediator of Gatad2a and Chd4 binding, and corresponds with previous publications which showed that the highly conserved Mbd2-CCR can form a heterodimer with Gatad2a and thus to modulate the complex’s function (Gnanapragasam et al., 2011).

### Interactions and assembly of Gatad2a-Mbd3/NuRD are context dependent and can be modified post-translationally

We and others have previously shown that the reprogramming factors (OSKM) factors directly interact and co-IP with Mbd3/NuRD complex following reprogramming initiation in somatic cells (van den Berg et al., 2010; Rais et al., 2013). By assessing these interactions in other cell states (representing different differentiation conditions), we noted that some of these interactions do not exist in naïve ESCs expanded in 2i/LIF conditions despite abundant expression of Oct4 and Klf4 (**Fig. 7A**) as well as expression of all known NuRD components under these conditions (**Fig. 7A**). However, upon brief 48h priming of naïve conditions and turning them into EpiLCs (**Fig. 7A**) or long term EpiSCs (**Fig. 7B**), establishment interactions with Oct4/Klf4 and Mbd3/NuRD became rapidly and strongly evident. These results are consistent with the notion that Mbd3/NuRD promotes ESC differentiation (Reynolds et al., 2012; Dos Santos et al., 2014; Yildirim et al., 2011), indicating that this propensity collates with increased NuRD-TF interactions upon pluripotency priming, or in somatic cell state conditions experiencing “non-physiologic” ectopic expression of pluripotency factors like OSK (**Fig. 7A-B**).

**Figure 7.**
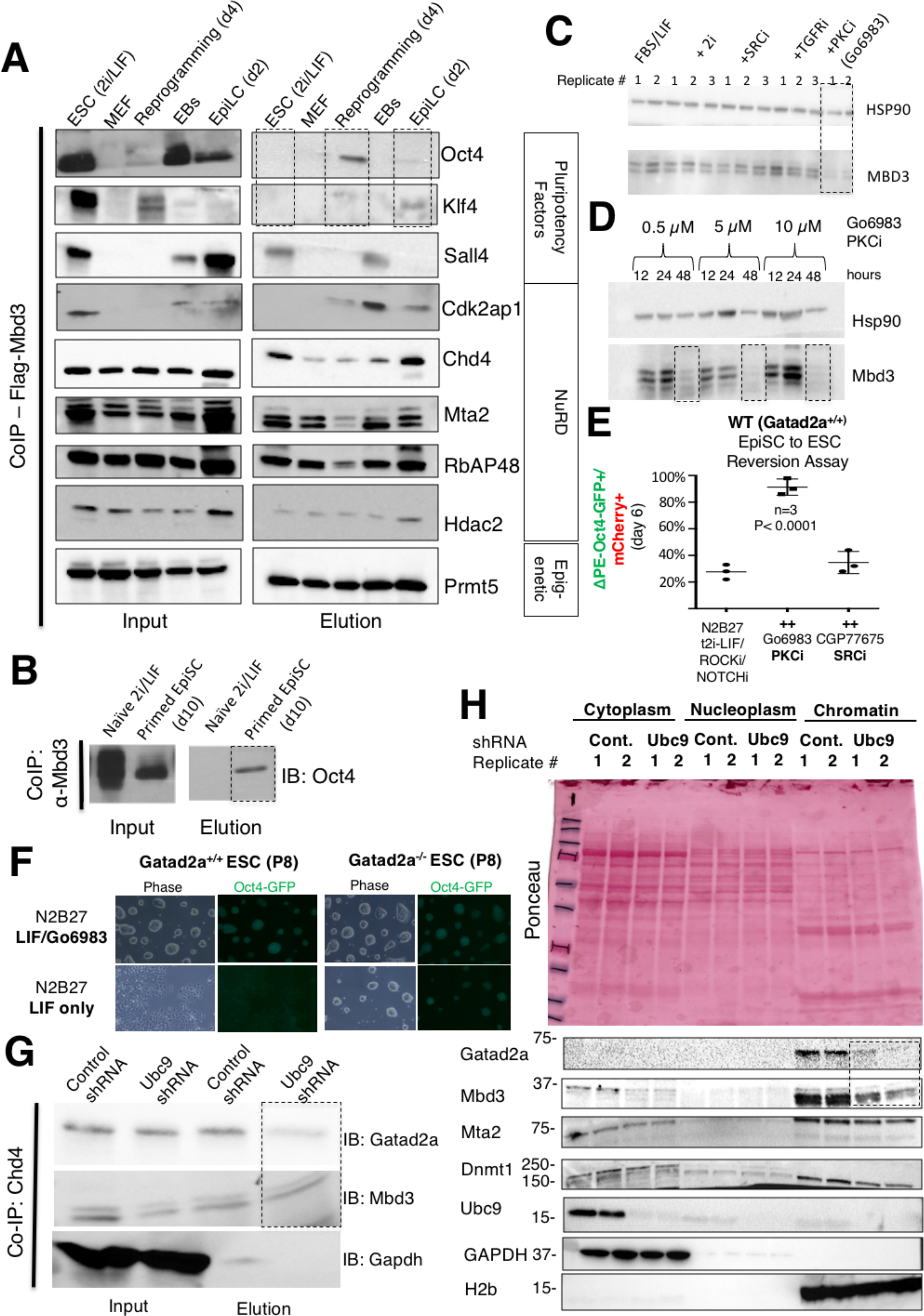
Interactions and assembly of Gatad2a-Mbd3/NuRD are context dependent and can be modified post-translationally. **A.** Rosa26-M2rtTA Col1a:TetO-2XFlag-Mbd3 ES cells were subjected to different differentiation protocols, and cells from 5 distinct states (naïve ESC, EpiLC, EBs, MEF, 4-day OSKM Reprogramming) were subsequently subjected to CoIP with anti- Flag-Mbd3. Lysates were then analyzed by western blot, and reacted with different antibodies against different NuRD components, Pluripotency factors, and other epigenetic proteins. While some of the proteins show constitutive binding to Mbd3 throughout all differentiation states (Mta2, Prmt5)- other proteins show differential binding (Oct4, Klf4, Cdk2ap1) and the latter is not correlated with the proteins level in the cell. **B.** Rosa26-M2rtTA Col1a:TetO-2XFlag-Mbd3 ES cells were either maintained in ground state naïve conditions or in priming conditions (Fgf2/Activin A). Lysates were subjected to Co-IP with anti-Flag-Mbd3 and examined by Western blot. Oct4 protein expression is significantly reduced after priming, but its binding to Mbd3 can be detected only in the primed pluripotent state. **C.** Mbd3 expression in ES cells treated with growth media containing different small molecules, after 72 hours of treatment. Unlike other treatments, PKCi Go6983 (5 μM) treatment resulted in a radical decrease in Mbd3 protein expression. **D.** PKCi Go6983 effect on Mbd3 level is seen after approximately 48 hours, in different concentration ranging from 0.5 to 10 μM. **E.** WT EpiSCs reversion efficiency to naïve ESCs in different conditions. Anova test P values are indicated. (one representative experiment out of 3 performed is shown). **F.** Isogenic WT and Gatad2a KO ESCs were expanded on feeder free gelatin coated plates in N2B27 LIF only or LIF/PKCi conditions. Phase images and Oct4-GFP signal maintenance are shown after 8 passages (P8). Oct4-GFP. **G.** ES cells treated with naïve ground state condition were treated with shRNA targeting Ubc9 or Scramble. Cells were lysed and subsequent CoIP of Chd4 shows a decrease in Gatad2a binding to the protein. **H.** Cells induced in naïve ground state 2i/LIF conditions and subsequent shRNA targeting either for Ubc9 or scramble negative control. The cells were lysed and fractioned – Cytoplasm, Nucleoplasm and Chromatin fractions, proteins were analyzed by western blot. The NuRD components Mbd3 and Gatad2a, but not Mta2, can be seen mainly in the chromatin fraction, and their expression is significantly reduced following shUbc9 treatment.

While the nature and molecular basis of these context specific interaction remain to be explored, we asked whether enhancing naïve pluripotency conditions by blocking other signaling pathways may deplete certain Mbd3-Chd4-Gatad2a/Complex components. Remarkably, of the several small molecule inhibitors previously published to promote naïve ESC maintenance (SRCi, TGFRi, PKCi) (Dutta et al., 2011; Han et al., 2010; Shimizu et al., 2012), the broad spectrum PKC inhibitor Go6983 dramatically depleted Mbd3 protein expression within 24 hours of treatment in different naïve conditions (**Fig. 7C-D**, **S8A-C**). The effect was not mediated by another PKC inhibitor, GF109203X (GFX), that does not target atypical PKC isoforms (**Fig. S8D**). Indeed, siRNA for the atypical PKCzeta isoform recapitulated partially the depletion seen in Mbd3 observed in ESCs (**Fig. S8E)**. This effect was seen in the context of pluripotent cells but not significantly when Go6983 was applied on a variety of somatic cells (**Fig. S8F**) suggesting that the Mbd3 depletion effect was not by specific and direct action of this small molecule on the NuRD complex (equivalent to example where ERKi depletes Uhrf1 and Dnmt3,a,b and l in mouse ESCs but not in somatic cells (Habibi et al., 2013)(Leitch et al., 2013). Finally, this reduction in Mbd3 protein was not accompanied by a change of Mbd3 mRNA transcript levels (**Fig. S8G**), suggesting rapid regulation at the post-translational level that will be of future interest for biochemical dissection.

We next tested the ability of Go6983 to boost iPSC efficacy formation from WT secondary reprogrammable MEFs. We observed an increase of reprogramming efficiency up to 45% compared to DMSO treated cells (**Fig S8H-I**). It is possible that the efficiencies were not as high to those seen in genetically modified MEF cells (**Fig. 2**), possibly because Go6983 does not deplete Mbd3 in somatic cells and only later after initiation of the process when cells start becoming ES-like (**Fig. S8 F, H**) (early stages of reprogramming when MEF identity is still maintained), which helps further boost reprogramming efficiency. However, in the context of EpiSC reversion, we were able to obtain up to 85% single cell EpiSC reversion efficiency in combination with LIF/ERKi/ROCKi/NOTCHi Vitamin C were supplemented with Go6983, while without PKCi or when using SRC instead, reversion efficiencies remained below ˜25% (**Fig. 7E**). Finally, while WT ESCs cannot maintain their naïve pluripotency in N2B27 LIF unless an additional component like 2i or PKCi are provided (Rajendran et al., 2013; Ying et al., 2003, 2008), Gatad2a-/- ESCs were fully stable for many passages in N2B27 LIF only conditions (in the absence of Go6983) as determined by the stringent naïve specific OG2 ΔPE-Oct4-GFP reporter (**Fig. 7F**). The latter results underscore a previously unidentified link between naïve PSC booster Go6983 and depletion of Mbd3/NuRD in the context of murine pluripotency and reprogramming.

Given the above result showing context- and signaling dependent influence on NuRD complex stability and function, we wondered whether some of other recently identified genetic perturbations shown to boost iPSC reprogramming efficiency had direct or indirect effects on NuRD assembly and its ability to be loaded on the chromatin. While we so far have not seen alteration in Mbd3/NuRD function following C/EBP induction in B cells (Di Stefano et al., 2016) or Dnmt1 inhibition in pre-iPSCs (Mikkelsen et al., 2008) (data not shown), we analyzed the effect of depleting global SUMOylation on iPSC reprogramming as it was shown to be an efficient booster (Borkent et al., 2016; Cheloufi et al., 2015). Moreover, we took special focus on SUMOylation since SUMOylation on Gatad2a protein in NIH3T3 cells **(Fig. S8J)** has been shown to be essential for its direct binding to Hdac1 within NuRD (Gong et al., 2006). Here, we identified a direct link between global inhibition of SUMOylation that boosts iPSC efficiency, to NuRD stability and assembly in the context of pluripotent cell reprogramming, specifically through Gatad2a-Chd4 interaction. We validated that knockdown of Ubc9 boosts WT MEF reprogramming up to ˜40% following OSKM induction as previously reported (**Fig S8I)**. The ability of Chd4 to Co-IP with Gatad2a was specifically decreased following Ubc9 knockdown in ES, but not in control shRNA used (**Fig. 7G**). Similar results were obtained when using a specific small molecule inhibitor (2-D08) to inhibit SUMOylation (**Fig. S8K**). Cellular fractionation, followed by chromatin isolation in control vs. Ubc9 depleted samples undergoing pluripotency reprogramming showed specific depletion of Gatad2a/Mbd3 from the purified chromatin fraction, but not DNMT1 or Mta2 (used as controls) (**Fig. 7H, Fig S8L**), supporting that Gatad2a-Mbd3/NuRD assembly and loading on the chromatin is compromised following depleting SUMOylation and may contribute, at least partially, to the increase of iPSC efficiency obtained under these conditions.

## Discussion

Over the last twelve years since the initial discovery of iPSCs, many approaches have been suggested for improving direct reprogramming efficiency and dynamics by the canonical Yamanaka factors (OSKM) (Takahashi and Yamanaka, 2016). Here we provide an additional example showing that radically efficient and deterministic direct induction of pluripotency is feasible with modified direct *in vitro* reprogramming approaches. The findings related to Gatad2a depletion and iPSC reprogramming characterized herein, expand work previously published by our lab that has demonstrated that optimized depletion of the NuRD complex component Mbd3, can alter this process and shift it toward deterministic dynamics and high efficiency (Rais et al., 2013). Nonetheless, complete depletion of Mbd3 during somatic state rapidly yields cell cycle malfunctions, thus technically restricting the usage of this platform to a carefully engineered set of mutant lines. In this work, we have further elucidated the mechanisms of Mbd3 repressive effect during reprogramming, and underlined the importance of NuRD complex recruitment, especially Gatad2a-Chd4. We have described Mbd3-Gatad2a-Chd4 as a triple component axis within NuRD complex (**Fig. S7F**), which mediates the inhibitory function of NuRD in the context of reprogramming and maintenance of naïve pluripotency. Disassembly of this sub-complex by different approaches, including elimination of one or more of its members, leads to efficient and (near-) deterministic reprogramming.

Results obtained herein underline the importance of the NuRD structure and sub-complexes; Chd4, Mbd3 and Gatad2a are all mutually exclusive in the NuRD complex and each can be replaced by a highly homologous protein (Chd3, Mbd2, Gatad2b, respectively). However, the specific and exact combination of all three seems to be crucial in order to execute the inhibitory function related to pluripotency and differentiation. This effect can be also seen in other physiological conditions (Bode et al., 2016) and development (e.g. brain cortical development) (Nitarska et al., 2016). Further, whether certain conformation of NuRD complex can also specifically enhances direct trans-differentiation between somatic cell types remains to be explored (Bussmann et al., 2009; Efe et al., 2011; Shu et al., 2013; Szabo et al., 2010; Vierbuchen et al., 2010).

While our results suggest that although the optimized depletion of Mbd3, Gatad2a or Chd4 result in a similar effect on iPSC reprogramming efficiency, these proteins hold differential roles, possibly in NuRD sub-complexes, or even in a NuRD-independent manner. This notion is supported by their different effect on cell proliferation and viability, and also expressed in their slightly different KO phenotypes during embryonic development, as both Chd4 and Mbd3 are embryonic lethal at a very early stage (E4.5-E6.5) (Kaji et al., 2007; O’Shaughnessy-Kirwan et al., 2015), while Gatad2a-KO embryos are viable until E8.5-10.5 (Marino and Nusse, 2007). Thus, along with the importance of the observations regarding the inhibitory role of Gatad2a-Mbd3 in the reprogramming process, it is clear that further work is needed to fully understand the mechanisms underlying Mbd3 functions (including NuRD dependent and independent ones) and specific role for Gatad2a, but not Gatad2b which can also assemble with Chd4/Mbd3. The findings that recruitment of OSKM by Mbd3/NuRD is prominent in primed or somatic state, but not in ground state naïve pluripotency, highlight the context dependent ability of NuRD interaction in different states whose molecular basis is of great future scientific interest.

Although this study shows in a variety of systems and assays that neutralizing Mbd3-Gatad2a-Chd4/NuRD complex facilitates up to 100% reprogramming efficiency, it remains to be seen whether all the cells follow the same molecular trajectory during the 8 days course or, alternatively, can achieve the final naïve state by distinct parallel paths. It is important to highlight that our results do not necessarily suggest that WT cells that succeed in generating iPSCs must neutralize this pathway (permanently or transiently) in order to succeed but may suggest that some cells succeed despite the negative effect of the presence of this complex. In other words, Gatad2aMbd3/NuRD may act like a negative rheostat rather than an absolute blocker for iPSC formation (Eggan, 2013; Smith et al., 2016; Zviran and Hanna, 2014), where it severely reduces the chance of conducive reactivation of correct reprogramming trajectory in Gatad2a/Mbd3 WT cells, since it represses that same genes OSKM are trying to reactivate (Rais et al., 2013). Upon dismantling this negative effector, nearly all cells embark correctly on a conducive reprogramming trajectory towards naïve pluripotency, which we characterize in detail in an accompanying manuscript by our group using the same platforms (accompanying submitted manuscript PDF copy is included with this submission) (Zviran et al., 2017).

A recent study has provided an NuRD independent example of the “gas and brakes” mechanisms for iPSC formation (Rais et al., 2013), where the OSKM reprogramming factors not only recruit complexes that positively promote reactivation of naïve pluripotency, but also recruit negative factors, in this case the NCoR/SMRT-HDAC3 co-repressor complex (Zhuang et al., 2018). The study showed up to 70% Oct4-GFP reactivation over a reprogramming period of 12 days by OSKM upon depletion of NCoR/SMRT (Zhuang et al., 2018)(in press – PDF attached with this submission). The latter is recruited directly by OSKM and negatively represses key loci required for successful reprogramming. The above results suggest that just like in the case of reprogramming factors that can act synergistically and occasionally substitute for each other to induce iPSCs (Festuccia et al., 2012; Martello et al., 2012; Qiu et al., 2015), multiple co-repressor complex like Mbd3/NURD, NCoR/SMRT, and potentially others may have overlapping effect in restraining the awakening of pluripotency is somatic cells (Rais et al., 2013; Zhuang et al., 2018).

Finally, it will be interesting to test whether by combining Gatad2a ablation with other manipulations that profoundly boost iPSC efficiency like transient C/EBPa expression in B cells (Di Stefano et al., 2013) or NCoR/SMRT depletion in MEFs (Zhuang et al., 2018), the kinetics and synchrony of the process can be further increased to reach that seen following somatic cell nuclear transfer where donor fibroblast turn on Oct4-GFP and acquire DNA global hypo-methylation by only up to 2 cell divisions (Gurdon, 2009; Jullien et al., 2014; Ma et al., 2014; Tachibana et al., 2013).

## Author Contributions

N.M and Y.R. conceived the idea for this project, designed and conducted experiments, and wrote the manuscript with contributions from all other authors. M.Z., R.M., A.A.C and Y.R. conducted micro-injections. S.B. and S.G. conducted and supervised high-throughput sequencing. A.Z. assisted in RNA-seq analysis. N.M., Y.L. and D. E. conducted MS analysis. W.J.G. and J.B. assisted in establishing and analyzing ATAC-seq experiments. E.C. assisted in DNA methylation sample preparation and analysis. N.N. supervised all bioinformatics analysis and analyzed bioinformatics data. S.P., S.H., T.S., J.B., O.G, M.A.E., L.L and V.K. assisted in tissue culture, preparing DNA reagents and reprogramming experiments. H.M. and Y.B. conducted and analyzed synchronous replication timing experiments. T.H. and S.G. assisted in Co-IP experiments. Y.R. and N.M engineered cell lines under S.V. supervision and designs. D.S. and Y.M. assisted and supervised experiments related to SUMOylation. N.N., Y.R. and J.H.H. supervised executions of experiments, adequate analysis of data and presentation of conclusions made in this project.

## Acknowledgements

We thank Bryce Carey, Yonatan Seltzer, Noam Stern-Ginossar and Naama Barkai for discussions. J.H.H is supported by a generous gift from Ilana and Pascal Mantoux, and research grants from the: European Research Council (ERC-Cog 2016 – CellNaivety), Flight Attendant Medical Research Council (FAMRI), Israel Science Foundation (ISF-ICORE, ISF-NFSC, ISF-INCPM … ISFMorasha programs), Kamin-Yeda program, Minerva fund, Israel Cancer Research Fund (ICRF), Human Frontiers Science Program (HFSP), the Benoziyo Endowment fund, New York Stem Cell Foundation (NYSCF), Kimmel Innovator Research Award, the Helen and Martin Kimmel Institute for Stem Cell Research. J.H.H. is a New York Stem Cell Foundation (NYSCF)–Robertson Investigator. N.N. is supported by ISF-Morasha program. We thank the Weizmann Institute management and board for providing critical financial and infrastructural support.

## Supplementary Figure Legends

**Supplementary Figure S1.**
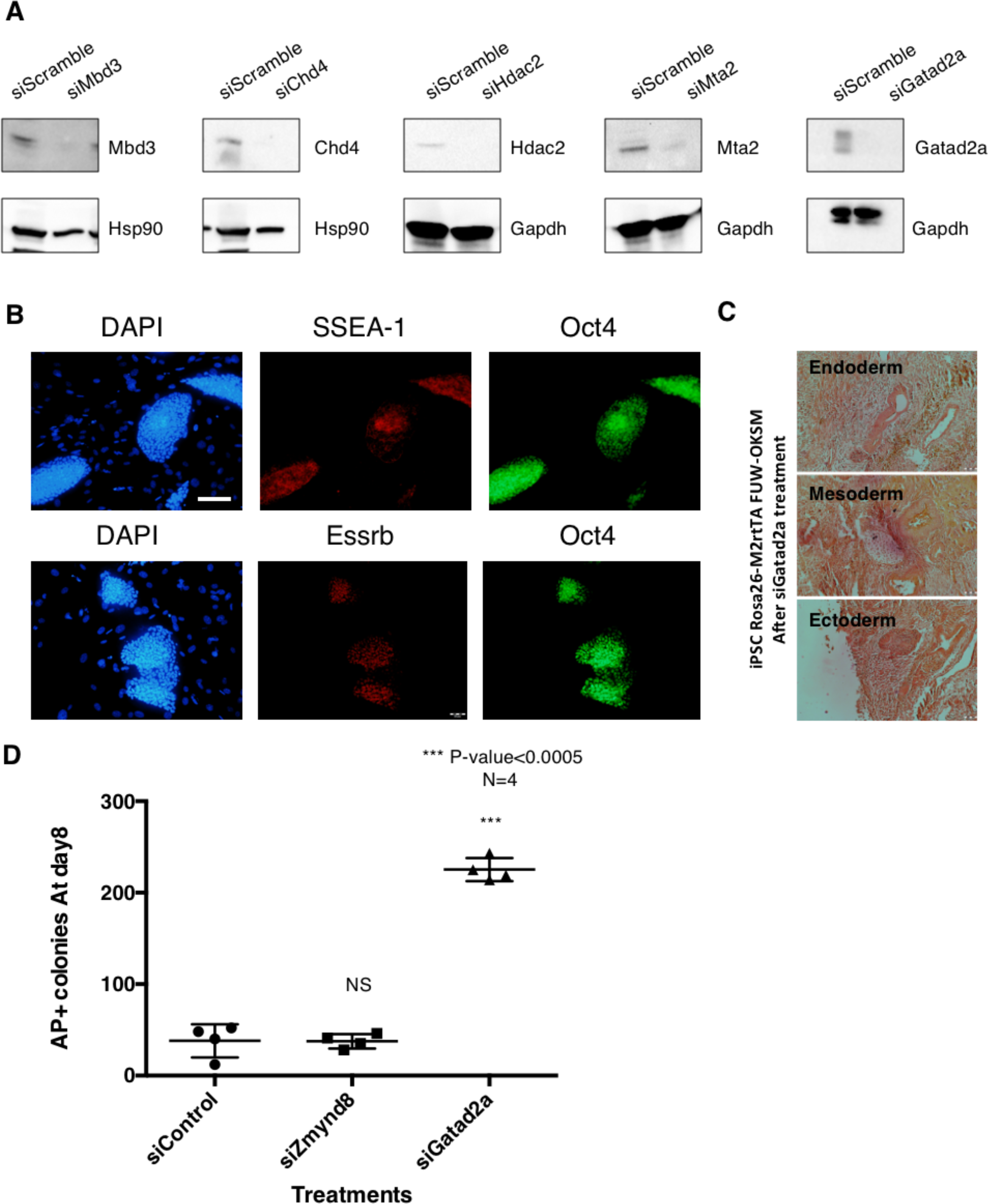
iPSC colonies formed following reprogramming with siRNA for Gatad2a are pluripotent. **A.** Western blot analysis following siRNA transfection for different NuRD components in MEFs, in order to validate specific protein deletion. **B.** Representative immunostaining of one of the iPSC clones derived from cells treated with siGatad2a (siGatad2a iPSC), stained positive to different pluripotent markers. Scale= 100 μM. **C.** *In vivo* teratoma formation following injection of an siGatad2a iPSC clone #2, demonstrating contribution to all three germ layers of development. D. MEFs harboring TetO-OKSM and M2rtTA cassettes were transfected with siRNA targeting Zmynd8 or Gatad2a, 2 and 4 days after reprogramming initiation following DOX administration. Reprogramming was than evaluated by AP staining at day 8 (n=4, two-sided Student’s t-test *p* values are indicated).

**Supplementary Figure S2.**
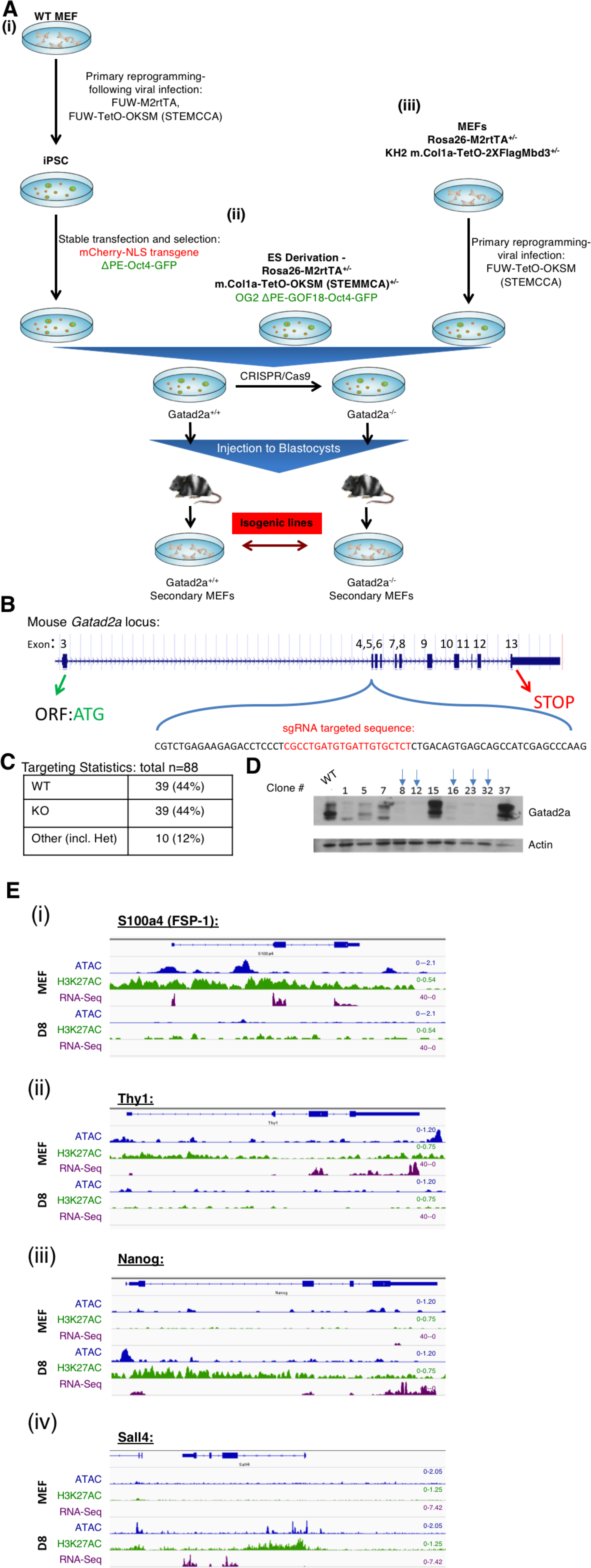
Generation of Gatad2a WT and depleted isogenic secondary platforms for iPSC reprograming. **A.** Three different clonal sets of isogenic secondary OSKM reprogrammable cells/sets were generated: (i) MEFs were reprogrammed to iPSC following viral infection of FUW-M2rtTA and FUW-OKSM. mCherry constitutive marker and ΔPE-Oct4-GFP markers were then introduced to the iPSC. (ii) R26-M2rtTA^+/-^ m. Col1a-OKSM^+/-^ ES were derived from E3.5 embryos following mating of mice. (iii) ES Kh2 m.Col1a-TetO-2XFlag-Mbd3 were injected to blastocysts, and MEF were harvested at E12.5. MEF were then reprogrammed following viral infection of FUW-OKSM. All described cell lines were then subjected to Gatad2a-KO using CRISPR/Cas9, followed by an injection of both the KO and its isogenic WT to blastocysts. Chimeric fibroblasts were separated from the donor cells by Puromycin selection. **B**. Summary of CRISPR/Cas9 strategy to generate Gatad2a null cells. sgRNA targeted sequence is indicated. **C**. Correct targeting efficiency by the strategy described in **B** to generate mouse Gatad2a KO PSCs as determined both by Western blot analysis and PCR sequencing. **D**. Representative Western blot analysis for sub cloned lines following targeting Gatad2a with sgRNA. Blue arrows indicate examples of KO clonal lines. **E.** Transcriptome landscape (RNA-seq), alongside ATAC-seq and H3K27Ac ChIP-seq, of fibroblasts (FSP-1 and Thy1) and pluripotent (Nanog and Sall4) related genes. IGV Data range is indicated at the top right corner of the signal.

**Supplementary Figure S3.**
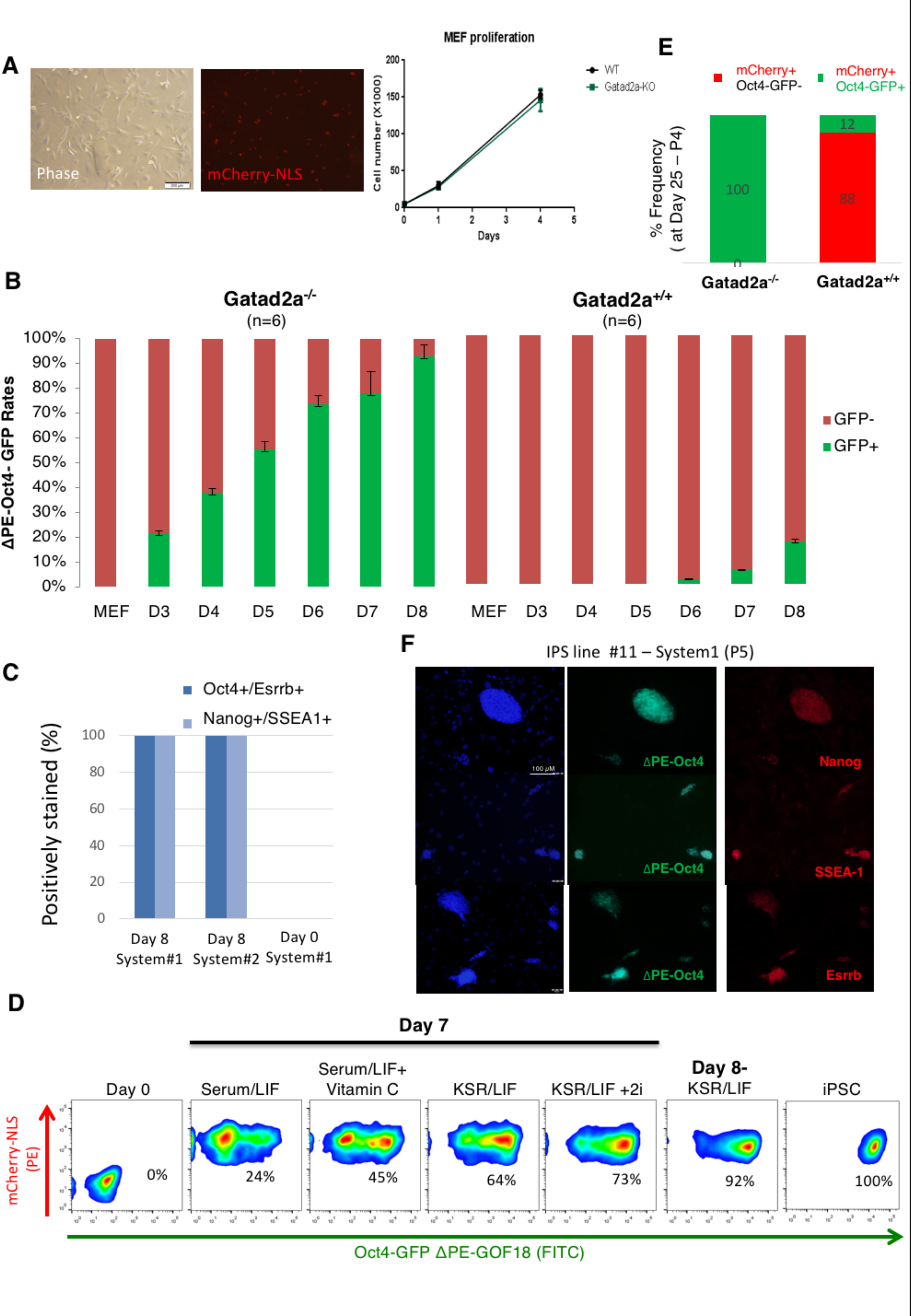
Gatad2a-KO has a dramatic beneficial effect on reprogramming efficiency in various conditions. **A.** Representative image of secondary Gatad2^-/-^ MEFs expanded in the absence of DOX. The cells had normal growth rate in comparison to their isogenic WT and passage matched Gatad2a^+/+^ MEF controls. **B.** A Comparison of ΔPE-Oct4-GFP reactivation dynamics’ statistics measured by flow-cytometry, in Gatad2a-WT and Gatad2a-KO isogenic platforms (n=6 wells per each time point). **C**. 96-well plates before and after 8 days of DOX induction were fixed and double stained for the indicated pluripotency markers. Frequency of positive scoring wells is indicated per each system and at different time points. **D.** Flow-cytometry measurements of ΔPE-Oct4-GFP reactivation in cells grown with different mediums. The beneficial effect of Gatad2a-KO was conserved in different mediums, including serum-based conditions. **E.** Monoclonal lines were established from Gatad2a^+/+^ and Gatad2a^-/-^ secondary cells and reprogrammed in 2i/LIF + DOX conditions as indicated in **Fig. 2A**. Fraction of Pre-iPSC partially reprogrammed lines (mCherry+/Oct4-GFP-) is indicated following measurement at Day 25 following reprogramming initiation (total of 4 passages). **F**. Representative immunostaining of a randomly selected clonal Gatad2a-KO iPSC (from system i) shows positive staining for all different bona fide mouse naïve pluripotency markers (scale bar = 100 μM).

**Supplementary Figure S4.**
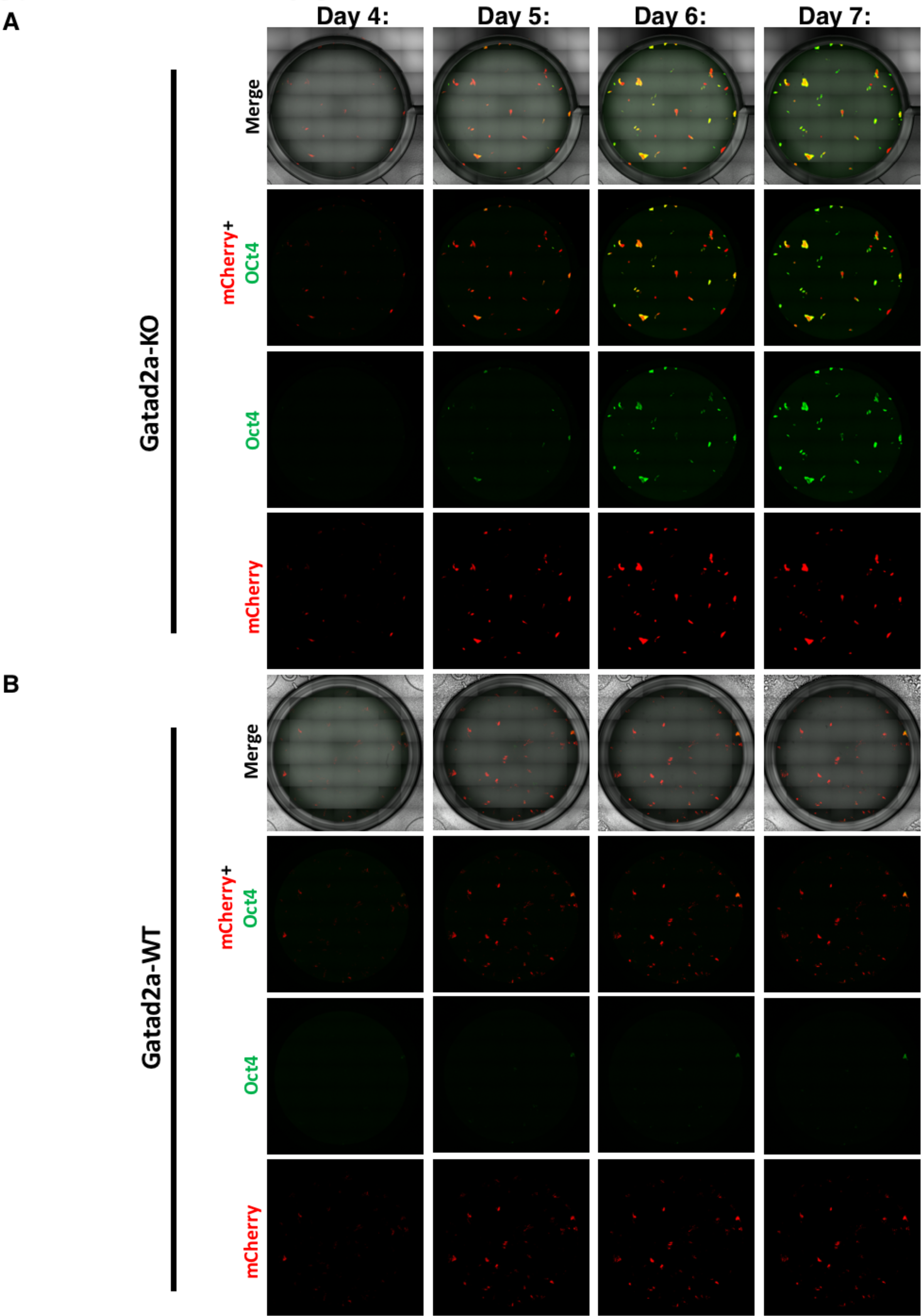
Full-well mosaic live-cell imaging of reprograming following Gatad2a ablation. Related to Fig. 3. Selected time-points from live-imaging of Gatad2a-KO and Gatad2a-WT reprogramming. Full-well mosaic of mCherry, Oct4-GFP and combined channels, shown for reprogramming of Gatad2a-KO (**A**) and its isogenic Gatad2a-WT paternal control (**B**) secondary MEF, harboring constitutive mCherry and ΔPE-Oct4-GFP reporter, at different time-points (days 3-7). Please also see **Supplementary Videos 1-2**.

**Supplementary Figure S5.**
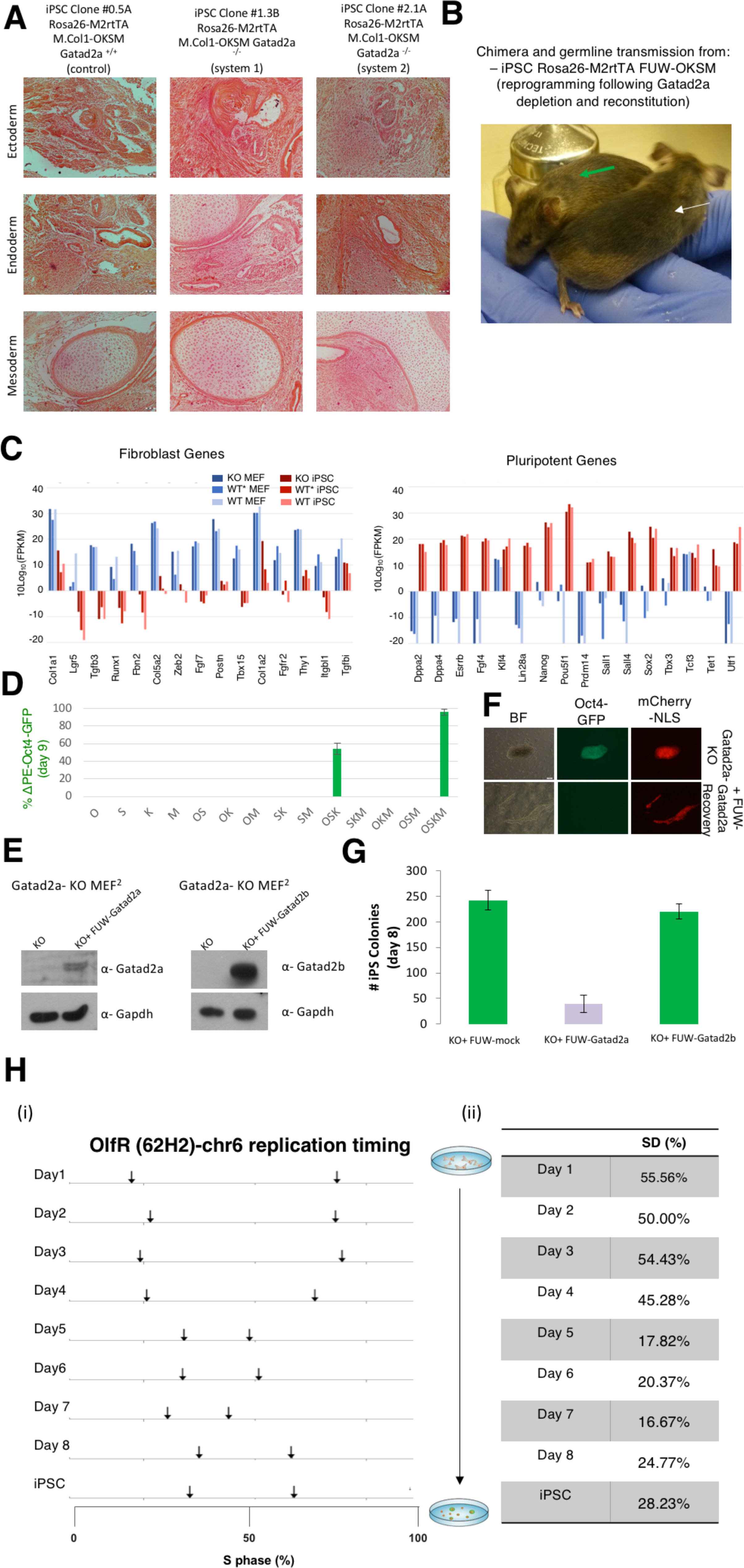
Characterization of iPSC lines obtained. **A.** Representative teratomas obtained following injection of the indicated iPSC lines. Mature differentiation was evident in all teratomas analyzed from all 15 randomly selected clones tested. **B.** A high contribution male chimeric mouse obtained following injection of Gatad2a-Tg rescued iPSC clone #1 (white arrow), and its agouti colored offspring (green arrow) obtained following mating with C/57B6 female, thus demonstrating successful germline transmission. **C.** Transcriptional level of selected fibroblast and pluripotent genes, as measured in MEF and iPS cells of 2 different systems: Gatad2a^-/-^, isogenic WT and non-isogenic WT* cells. **D.** Reprogramming efficiency of Gatad2a^-/-^ MEFs after transduction with the indicated combinations of reprogramming factors at day 9. Polycistronic vectors were used for OSK and OSKM combinations. One representative experiment of three independent sets is shown. **E.** Protein validation of Gatad2a or Gatad2b transgenic (Tg) over-expression following lentiviral transfection in Gatad2a-KO MEF. **F.** Representative image shows how partially reprogrammed Pre-iPSCs can be obtained only from lines with transgenic rescued Gatad2a expression (mCherry+/ΔPE-Oct4-GFP-; lower panels), but not from parental Gatad2a-KO cells (only mCherry+/#PE-Oct4-GFP+ cells can be obtained; upper panels). **G.** Overexpression of Gatad2a, but no Gatad2b, in Gatad2a-KO reprogrammable MEF reduces reprogramming efficiency comparing to mock overexpression control (N=4 per sample). **H.** Gatad2a-KO secondary MEF were subjected to reprogramming. At each time-point, the cells were labeled with BrdU, and replication time was examined using OlfR probe (Chromosome 6), as described (Masika et al., 2017). Approximate replication time of each allele is graphically described in (i) and in table (ii) summarizes replication time (for different time points days 1-8 and iPSC number of cells analyzed: n= 90,106, 79, 106, 101, 108, 42, 109, 124).

**Supplementary Figure S6.**
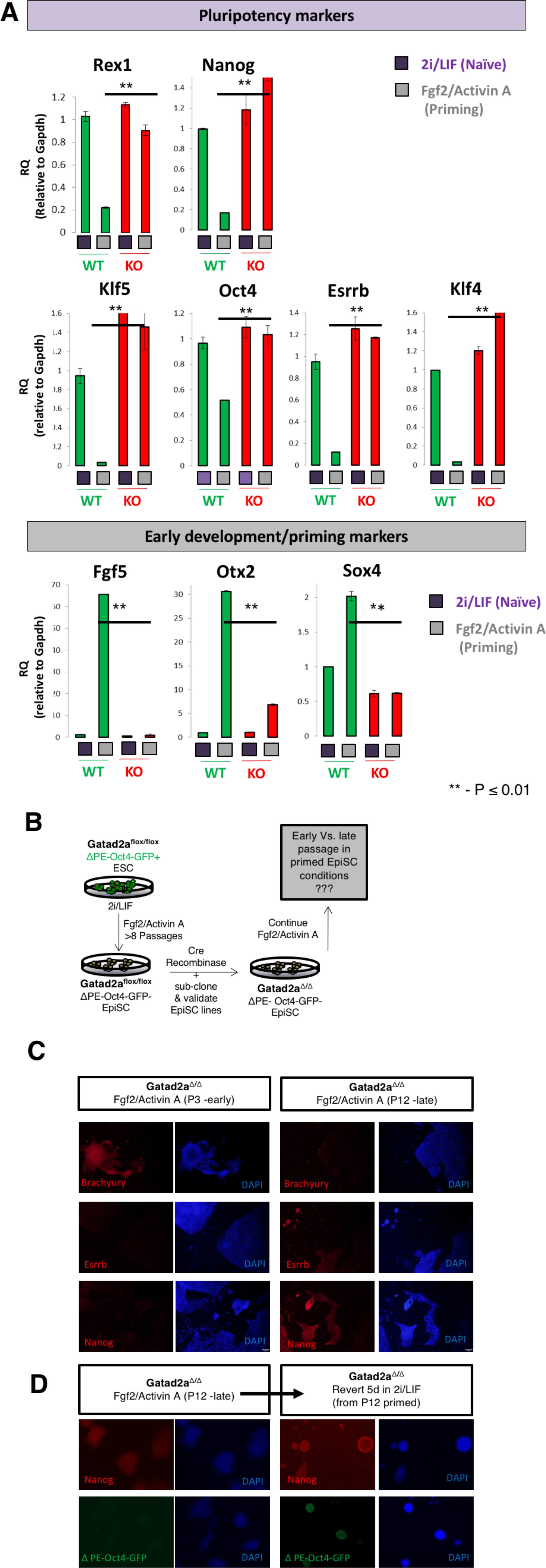
Gatad2a ablation compromises induction and maintenance of primed murine pluripotency. **A.** RT-PCR analysis of different pluripotency and early differentiation markers before and after Fgf2/Activin A exposure, in isogenic Gatad2a^+/+^ and Gatad2a^-/-^ ESCs. **B**. Gatad2a^flox/flox^ naïve ES were subjected to priming using FGF/Activin A for 8 passages, followed by Cre-treatment in order to delete Gatad2a and sub clone isogenic Gatad2a^Δ^/Δ EpiSC lines. Subsequently, Gatad2a^Δ^/Δ EpiSC were characterized at early and late passages in EpiSC conditions. **C.** Immunostaining analysis showed that with extended passaging Gatad2a^Δ^/# began to re-activate naïve pluripotency markers (Esrrb and Nanog) and to down-regulate differentiation markers, such as Brachyury despite the absence of 2i/LIF conditions and continued expansion in Fgf2/Activin A primed conditions. **D.** Nanog and ΔPE-Oct4-GFP profiled in late passage Gatad2a^Δ^/Δ EpiSCs and after 5 days reversion in 2i/LIF.

**Supplementary Figure S7.**
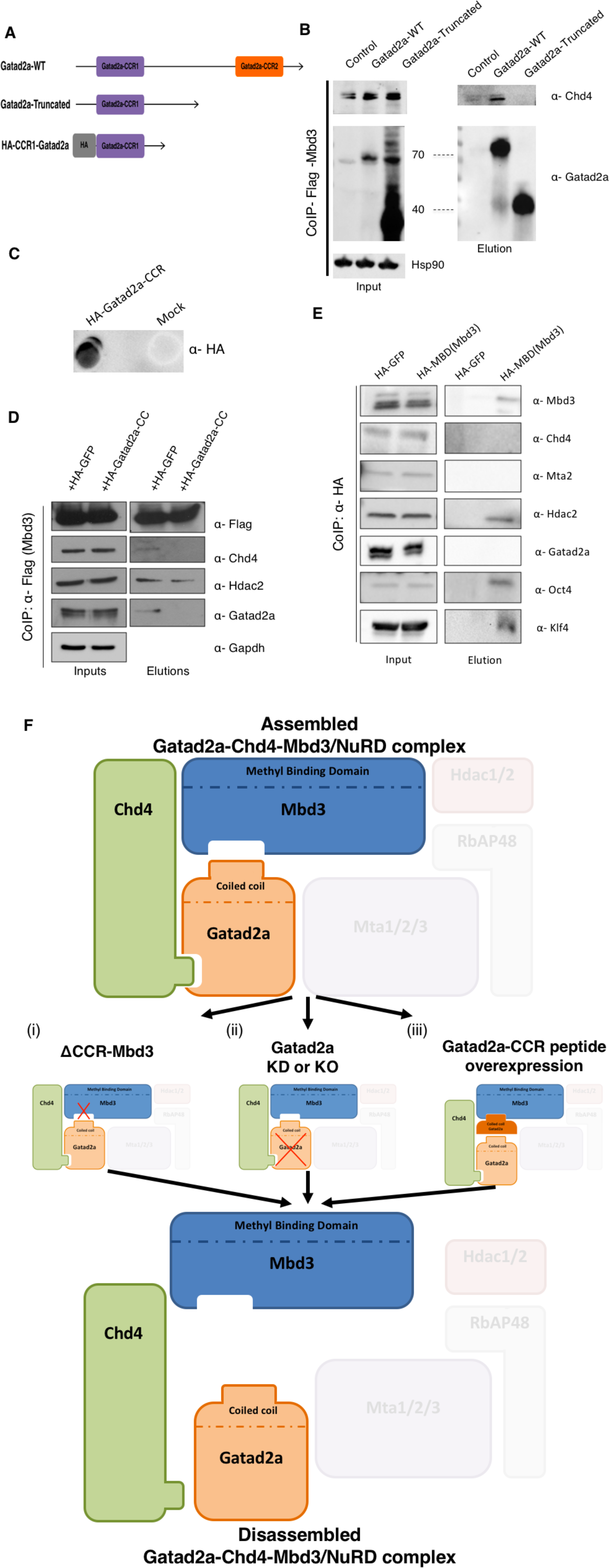
Gatad2a, Mbd3 and Chd4 constitute a critical axis within the NuRD complex mediating iPSC reprogramming inhibition. **A.** A scheme of Gatad2a constructs and mutants used. Gatad2a-WT, Gatad2a-Truncated (lacking the CCR2 of Gatad2a) and HA-tagged-CCR1-Gatad2a construct. **B.** Flag-Mbd3 and different Gatad2a constructs were cotransfected in 293T cells. Flag-Mbd3 binds both forms of Gatad2a (WT and truncated), as shown by Flag-Mbd3 CoIP. **C.** Dot blot analysis to validate protein expression following HA-Gatad2aCCR1 peptide overexpression. **D.** Overexpression of Flag-Mbd3 with HA-Gatad2a-CCR1 or control (HA-GFP) in 293T cells, followed by Co-IP for anti-Flag-Mbd3. Over-expression of Gatad2a-CCR1 reduces the binding of Mbd3 to endogenous Gatad2a and Chd4, comparing to the control specimen. Further, CCR can cause a reduction in Mbd3 binding to endogenous Gatad2a and Chd4, without changing other NuRD components (such as Hdac2) binding. **E.** Overexpression of STEMCCA-OKSM vector, HA-MBD (methyl-binding domain of Mbd3) or HA-GFP in 293T cells, was followed by Co-IP with anti-HA. The results demonstrate that MBD domain can bind to pluripotency factors such as Oct4 and to the NuRD component Hdac2 but cannot bind Gatad2a or Chd4. **F.** Summarizing scheme for three different approaches for neutralizing Mbd3-Gatad2a-Chd4 axis: (i) by deleting the CCR domain of Mbd3 (ii) KO or KD of Gatad2a protein and (iii) overexpression of an exogenous Gatad2a-CCR competitive peptide that interferes with Gatad2aCCR interaction with Mbd3.

**Supplementary Figure S8.**
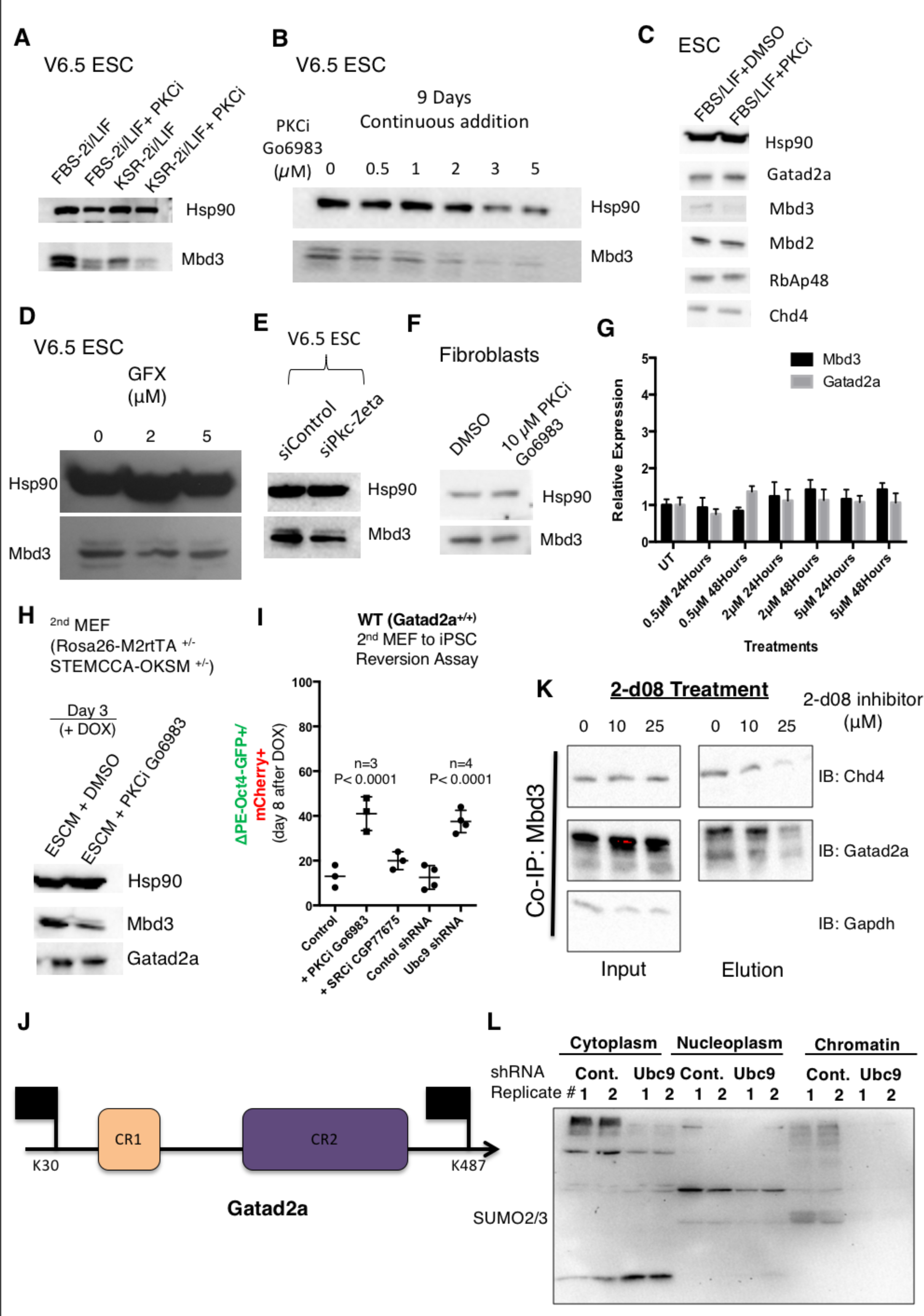
Interactions and assembly of Gatad2a-Mbd3/NuRD are context dependent and can be modified post-translationally. **A.** Western blot showing Mbd3 protein levels in ESCs expanded in the indicated growth conditions. Mbd3 levels were decreased following PKCi (Go6983) treatment in ES cells maintained in different conditions (FBS and KSR based growth media, supplemented with 2i). **B.** Reduction in Mbd3 protein expression is maintained following long term PKCi (Go6983) treatment, as examined in different concentrations (0.5-5 μM). Media was exchanged ever 48h. **C.** Western blot analysis for different NuRD components in ESC cells with and without Go6983. Analysis shows that other NuRD components expression level is not affected from PKCi (Go6983) treatment. **D.** as in A, but a different PKCi was used, termed GFX which does not inhibit atypical PKC pathway. **E.** Mbd3 protein levels following shRNA treatment of mouse V6.5 ESCs for atypical PKCzeta isoform. **F.** Mbd3 levels in MEFs following treatment with Go6983 are not affected. **G.** RT-PCR analysis for Mbd3 transcript abundance 24 and 48h following PKCi Go6983 treatment, do not show significant changes and reduction in Mbd3 transcript level. This suggests that the depletion in Mbd3 protein levels shown in **A-C** is post-translational. **H.** Go6983 causes mild depletion in Mbd3 in MEFs following 3 days of OSKM expression. **I**. Gatad2a^+/+^ WT MEFs carrying ΔPE -Oct4-GFP reporter and constitutive mCherry markers (secondary system i) were subjected to iPSC reprogramming protocols in in **Fig. 2A** and iPSC efficiency was quantified at day 8. In the last two conditions included in the panel, the MEFs were pre-treated with control and Ubc9 shRNA following with Neomycin selection (Cheloufi et al., 2015), and then subjected to DOX mediated iPSC reprogramming. Anova P values are indicated. **J.** A Schematic representation of Gatad2a known domains and confirmed SUMOylation sites (Gong et al., 2006). Importantly, harnessing prediction tools (Zhao et al., 2014) for identification of possible SUMO consensus sites results in 2 additional sites inside the coiled coil regions (Chi-squared test *p* value<0.05). **K.** 2-d08 specific small molecule inhibitor for SUMOylation was applied during co-IP experiments for Chd4 at the indicated increasing concentrations known to deplete global SUMOylation levels. Cells were lysed the CoIP of Mbd3 shows a decrease in Gatad2a binding to Chd4, as similarly seen with shRNA depletion of Ubc9 (**Fig. 7G**). **L.** As in **Fig. 7H** with immunoblot for SUMO2/3 on the same exact gel series.

**Supplementary Video 1 – Whole-well unbiased mosaic live-cell imaging of isogenic Gatad2aWT and Gatad2a-KO cells during reprogramming.** Two movies presented side by side (Left- Gatad2a-WT, Right- Gatad2a-KO), acquired as live-cell imaging of isogenic secondary MEF, harboring constitutive mCherry-NLS and ΔPE-Oct4-GFP reporter, during OKSM mediated reprogramming. The two lines are isogenic and differ only by the ablation of Gatad2a. The movie describes days 3-7 of the process, as can be seen in 4 different panels- including mCherry only (showing only the proliferation of the viable donor cells) and GFP only (showing naïve state induction efficiency).

**Supplementary Video 2 – Whole-well unbiased mosaic live-cell imaging of Gatad2a-KO during iPSC reprogramming.** Live-cell imaging of Gatad2a-KO secondary MEF, harboring constitutive mCherry-NLS and #PE-Oct4-GFP reporter, during OKSM mediated reprogramming. The movie describes 4 different replicates during days 3-7 of the reprogramming process, maintained in the same conditions.

## Methods

### Mouse stem cell lines and cell culture

WT or Mutant mouse ESC/iPSC lines and sub-clones were routinely expanded in mouse ES medium (mESM) consisting of: 500ml DMEM-high glucose (ThermoScientific), 15% USDA certified Fetal Bovine Serum (Biological Industries), 1mM L-Glutamine (Biological Industries), 1% nonessential amino acids (Biological Industries), 0.1mM β - mercaptoethanol (Sigma), penicillin-streptomycin (Biological Industries), 10μg recombinant human LIF (Peprotech). For ground state naïve conditions (N2B27 2i/LIF), murine naïve pluripotent cells (iPSCs and ESCs) were conducted in serum-free chemically defined N2B27-based media: N2B27-based media: 250ml Neurobasal (ThermoScientific), 250ml DMEM:F12 (ThermoScientific) 5ml N2 supplement (Invitrogen; 17502048), 5ml B27 supplement (Invitrogen; 17504044), 1mM glutamine (Invitrogen), 1% nonessential amino acids (Invitrogen), 0.1mM β -mercaptoethanol (Sigma), penicillin-streptomycin (Invitrogen), 5mg/ml BSA (Sigma), small-molecule inhibitors CHIR99021 (CH, 3 $M- Axon Medchem) and PD0325901 (PD, 1 $M - Axon Medchem). Primed N2B27 media for murine cells (EpiSCs or EpiLCs) contained 12ng/ml recombinant human FGF2 (Peprotech Asia) and 20ng/ml recombinant human ACTIVIN A (Peprotech) (instead of 2i/LIF). Mycoplasma detection tests were conducted routinely every month with MycoALERT ELISA based kit (Lonza) to exclude mycoplasma free conditions and cells throughout the study.

### Conversion of naïve ESC to EpiLC or EpiSC

Naïve ESCs were seeded on Matrigel coated plates, at 5X10^5^ cells per well (6-wells plate), in naïve N2B27 2i/LIF conditions after being contained in those conditions for at least 10 days. After 24 hours, medium was changed to N2B27 supplemented with FGF2 (12ng/ml, Peprotech) and ACTIVIN A (20ng/ml, Peprotech). EpiLCs were harvested and validated at 48-96 post induction of priming. EpiSCs, which are lines stably expanded in FGF2/ACTIVIN A conditions were maintained on Matrigel coated plates and passaged with collagenase every 4-5 days as previously described (Brons et al., 2007; Geula et al., 2015; Tesar et al., 2007).

### Knockout by CRISPR/Cas9 targeting

Different PSC lines (as elaborated previously) were genetically manipulated by CRISPR/Cas9, in order to achieve Gatad2a-KO on different genetic backgrounds. Cells were transfected (Xfect, clontech) with Cas9-sgRNA Gatad2a (CGCCTGATGTGATTGTGCTCT) on px330 backbone, alongside mCherry-NLS plasmids. The cells were sorted after 72 hours (mCherry positive cells) under sterile conditions via ND FACS ARIA III, and seeded on irradiated MEFs. After 8 days, colonies were picked and examined by HRM analysis (MeltDoctor HRM master mix, Life technologies, #4415440). Candidate colonies were analyzed by western blot and confirmed by sequencing. For generation of floxed conditional knockout Gatad2a cells, CRISPR/Cas9 and donor vector were used as described (**Fig 4E**). Briefly, cells were transfected by electroporation with both donor vector containing Gatad2a Exon2 flanked by lox sequences, and px330 Cas9-sgRNA (gatacccatggtgcccacgg). Cells were subjected to selection by G418 and subsequent analysis of selected clones by Southern Blot. Southern Blot analysis was used to confirm correct targeting. 10-15Mg of genomic DNA was digested with HindIII restriction enzyme for 5 hours and separated by gel electrophoresis. The DNA was transferred to a nitrocellulose membrane that was next hybridized with a radioactive labeled probe and developed using ECL (ThermoScientific). Positive clones were transfected with Flippase to excise the neomycin selection cassette.

### Secondary reprogrammable cells and systems

Please see **Fig. S2A** for schematics summarizing the three isogenic systems detailed below:

<u>System i: FUW-M2RtTA; FUW-TetO-STEMCCA, ΔPE-GOF18-Oct4-GFP cells:</u> 129*Jae* MEFs were transduced with constitutively expressed FUW-M2RtTA and with FUW-TetO-STEMCCAOKSM (Sommer et al., 2009) via standard lentiviral preparation and infection. Cells were then reprogrammed in mESM + DOX for 14 days, and number of clonal iPSC line were established. A randomly selected clone was then transduced with ΔPE-Oct4-GFP carrying Zeomycin resistance (Addgene # 52382) (Gafni et al., 2013; Rais et al., 2013). Randomly selected clone validated for specific activity of the ΔPE-GOF18-Oct4-GFP reporter (downregulated upon priming and completely shut off upon differentiation 10 day EBs), was next labeled with a constitutively expressed mCherry-NLS cassettes using electroporation (Addgene #52409) (Rais et al., 2013). Only then, a sub cloned clone was subjected to CRISPR/Cas9 targeting in order to generate and isogenic Gatad2a-KO cell line which was used as a source for secondary somatic cells or EpiSCs and used for conversion.

<u>System ii: M.Col1a:STEMCCA-OKSM^+\-^; Rosa26-M2rtTA; ΔPE-GOF18-Oct4-GFP cells:</u> R26-M2RtTA homozygous mice (Jackson #006965) were mated with m.Col1a-TetO-STEMCCAOKSM (Jackson #011001, kindly provided by K. Hochedlinger) homozygous mice, in order to produce M2rtTA-OKSM mice double positive mice. The F1 mice were mated in order to create double homozygous offspring (RTTA ^+\+^ OKSM ^+\+^). F2 offspring (Double homozygous rtTA^+\+^ OKSM^+\+^) were mated with previously generated ΔPE-GOF18-Oct4-GFP homozygous reporter transgenic mice (also known as OG2 strain and line) (Jackson 004654, kind gift by H. Scholer), in order to generate a double heterozygote with Oct4 reporter offspring (M2rtTA ^+\-^ OKSM ^+\-^ ΔPEGOF18-Oct4-GFP ^+\-^. Subsequently, mouse ESCs were derived in mESM-MEF conditions and a cell line carrying the above genotype was transduced with a constitutively expressed mCherry-NLS cassettes using electroporation. Only then, a sub cloned clone was subjected to CRISPR/Cas9 targeting in order to generate and isogenic Gatad2a-KO cell line which was used as a source for secondary somatic cells or EpiSCs and used for conversion.

<u>System iii: FUW-TetO-STEMCCA-OKSM R26-rtTA Cola:tetO-3XFlag-Mbd3:</u> MEFs generated via microinjection of R26-rtTA col1a:TetO-3XFlag-Mbd3 ESCs, were introduced with OKSM (FUW-TetO-STEMCCA-OKSM) using a lentiviral transduction. An iPSC cell line was established in mESM + DOX conditions, and subsequent Gatad2a-KO was achieved using CRISPR-Cas9 genome editing. In all of the above, both the Gatad2a-KO and the isogenic Gatad2a-WT were then injected to a blastocyst, and MEF were harvested at E12.5-E13.5. Chimeric MEF were then isolated using Puromycin resistance.

Secondary mouse embryonic fibroblast (MEF) from Mbd3^flox/-^ cell line (A12 clone: *Mbd3^flox/^−* cell lines that carries the GOF18-Oct4-GFP transgenic reporter (complete *Oct4* enhancer region with distal and proximal enhancer elements) (Addgene plasmid #60527)) and WT* cell line (WT-1 clone that carries the deltaPE-GOF18-Oct4-GFP reporter (Addgene plasmid#52382) were previously described (Rais et al., 2013). Note that we do not use Oct4–GFP or any other selection for cells before harvesting samples for conducting genomic experiments. All animal studies were conducted according to the guideline and following approval by the Weizmann Institute IACUC (approval # 33550117-2 and 33520117-3). Cell sorting and FACS analysis were conducted on 4 lasers equipped FACS Aria III cells sorter (BD). Analysis was conducted with either DIVA software or Flowjo.

### Deterministic Fibroblast to iPSC Reprogramming experiments

Deterministic or near-deterministic iPSC reprogramming experiments were executed according to the following protocol, unless stated otherwise; Reprogramming was initiated in mESM, which contained 500 ml DMEM-high glucose (ThermoScientific), 15% USDA certified fetal calf serum (Biological Industries), 1 mM L-glutamine (BI), 1% non-essential amino acids (BI), 1% penicillin– streptomycin (BI), 0.1 mM β -mercaptoethanol (Sigma), 10 μg recombinant human LIF (Peprotech). MES medium for reprogramming was supplemented with Doxycycline (1 μg/ml), and the MEF^2^ were seeded on irradiated DR4 feeders. After 3.5 days, the medium was changed to serum-free media: referred to as KSR-2i/LIF media, which was also supplemented with Doxycycline, as well as 2i/LIF. The medium contained- 500 ml DMEM – high glucose (ThermoScientific), 15% Knockout Serum Replacement (KSR - ThermoScientific; 10828), 1 mM L-Glutamine (BI), 1% non-essential amino acids (BI), 0.1 mM β-mercaptoethanol (Sigma), 1% penicillin–streptomycin (BI), 10 μg recombinant human LIF (Peprotech), CHIR99021 (3 μM; Axon Medchem), PD0325901 (PD, 1μM; Axon Medchem). Cells were seeded upon irradiated MEF on 0.2% gelatin coated plates, at a low density of approximately 100-150 cells/cm. Reprogramming was conducted at 5% O2 5% Co2 and 37C in the first 2.5 days, and then moved to 20% O2 5%Co2 and 37C. Media replaced every other day (0, 2, 3.5, 6, 8). No blinding was conducted when testing outcome of reprogramming experiments throughout the study. For all mouse iPSC reprogramming experiments intended for genomics analysis, irradiated human foreskin fibroblasts were used as feeder cells (rather than mouse DR4 MEFs), as any sequencing input originating from the use of human feeder cells cannot be aligned to the mouse genome and is therefore omitted from the analysis.

### Knockdown by siRNA transfection

siRNA transfections were carried using Lipofectamine RNAiMAX (ThermoScientific #13778075), according to manufacturer protocol. Briefly, cells were seeded 24 hours before the transfection took place, in order to achieve approximately 70% confluent adherent cells. Next, both siRNA and Lipofectamine were diluted in Opti-MEM, in accordance to plate size (for a 6-wells plate well, 20pmole of siRNA were applied). Both the reagent and siRNA were mixed and vortexed, then left on RT for 5 minutes; the mixture was subsequently added to cells. Cell growth Media was changed after 24 hours, and cells for protein samples were harvested 48 hours after transfections. When siRNA transfection was a part of a reprogramming experiment, cells were retransfected every 48 hours, in order to maintain the knockdown, unless stated otherwise. Knockdown was confirmed by Western blot analysis.

The siRNA used are the following-

|  |  |  |
| --- | --- | --- |
| Mouse-Mbd3 | Invitrogen | MSS-237238 |
| Mouse-Chd3 | Invitrogen | MSS238382, MSS238383, MSS238384 |
| Mouse-Chd4 | Invitrogen | MSS- 200894, MSS-200895, MSS-200896 |
| Mouse-Gatad2a | Invitrogen | MSS239240, MSS239241, MSS239242 |
| Mouse-Zmynd8 | Invitrogen | MSS213659, MSS213660, MSS213661 |
| Mouse- Gatad2b | Invitrogen | MSS213785, MSS213786, MSS213787 |
| Mosuse-Mta2 | Invitrogen | MSS215840, MSS215841, MSS215842 |
| Mouse-Hdac2 | Invitrogen | MSS205071, MSS205072, MSS205073 |
| Mouse-Mbd2 | Invitrogen | MSS206540, MSS206541, MSS206542 |
| Mouse-Prkcz | Invitrogen | S71714 |
| Stealth RNAi™<br>siRNA Scrambled<br>Negative Control,<br>Med GC | Invitrogen | 12935300 |

### DNA plasmids and mutants

Additional DNA plasmids used in this study for ectopic expression included: Fuw-Flag-Mbd3 (Addgene 52371), FUW-Flag-Mbd3 delta1-70 (Addgene 52372). Newly generated plasmids include: FUW-2XFlag-ΔCCR-Mbd3 (Deletion of 220-279), FUW-Gatad2a, FUW-Gatad2b, pCAG-HA-Gatad2a-CCR (140-174 of Gatad2a), pCAG-HA-(MBD)Mbd3 (1-170 of Mbd3), pCAG-HA-GFP. All new plasmids will be made available through Addgene following publication of this manuscript.

### Western blot and dot blot analysis

Cells were harvested, and whole cell protein was extracted by lysis buffer, containing 150 mM NaCl, 150 mM Tris-Hcl (PH=7.4), 0.5% NP40, 1.5 mM MgCl2, 10% Glycerol. Protein’s concentration was determined by BCA Kit (ThermoScientific). Blots were incubated in the following primary antibodies (diluted in 5% BSA in PBST):

|  |  |  |
| --- | --- | --- |
| Mbd3 | 1:1000 | Bethyl A302-529A |
| Mbd2 | 1:1000 | Bethyl A301-633A |
| Gatad2a (H-162) | 1:1000 | Santa Cruz Sc-134712 |
| Gatad2b | 1:1000 | Bethyl A301-281A |
| Hdac2 (c-8) | 1:1000 | Santa Cruz sc-9959 |
| Mta2 (c-20) | 1:500 | Santa Cruz sc-9447 |
| Chd4 | 1:1000 | Abcam Ab70469 |
| Oct4 (H-134) | 1:1000 | Santa Cruz sc-9081 |
| Klf4 (H-180) | 1:1000 | Santa Cruz Sc20691 |
| Nanog | 1:1000 | Behtyl, A300-398A |
| Flag-M2 | 1:1000 | Sigma F3165 |
| Ha.11 16B12 | 1:1000 | Covance MMS-101R |
| Gapdh | 1:5000 | Epitomics 2251-1 |
| Hsp90β | 1:1000 | Epitomics 1492-1 |
| Prmt5 | 1:1000 | Millipore 07-405 |
| Cdk2ap1 (H-4) | 1:500 | Santa Cruz sc-390283 |
| RbAp48 (N-19) | 1:1000 | Santa Cruz sc-8270 |
| Sall4 | 1:1000 | Abcam ab29112 |
| Ubc9 | 1:1000 | Cell signaling #4918 |
| Dnmt1 | 1:1000 | Abcam ab87654 |

Secondary antibodies used: Peroxide-conjugated AffiniPure goat anti-rabbit (1:10,000, 111-035-003; Jackson ImmunoResearch). Blots were developed using SuperSignal West Pico Chemiluminescent substrate (ThermoScientific, #34080). Dot-blot method was used to identify small peptides expression. Briefly, protein lysate was loaded to a nitrocellulose membrane and was left to air-dry. Membrane was then blocked by 5% milk, and later incubated with the antibody.

### Real Time (RT)-PCR analysis

Total RNA was extracted from the cells using Trizol. 1 μg of RNA was then reverse transcribed using high-capacity cDNA reverse Transcription kit (Applied Biosystems). Quantitative PCR was performed with 10ng of cDNA, in triplicates, on Viia7 platform (Applied Biosystems). Error bars indicate standard deviation of triplicate measurements for each measurement. The primers in use were the following:

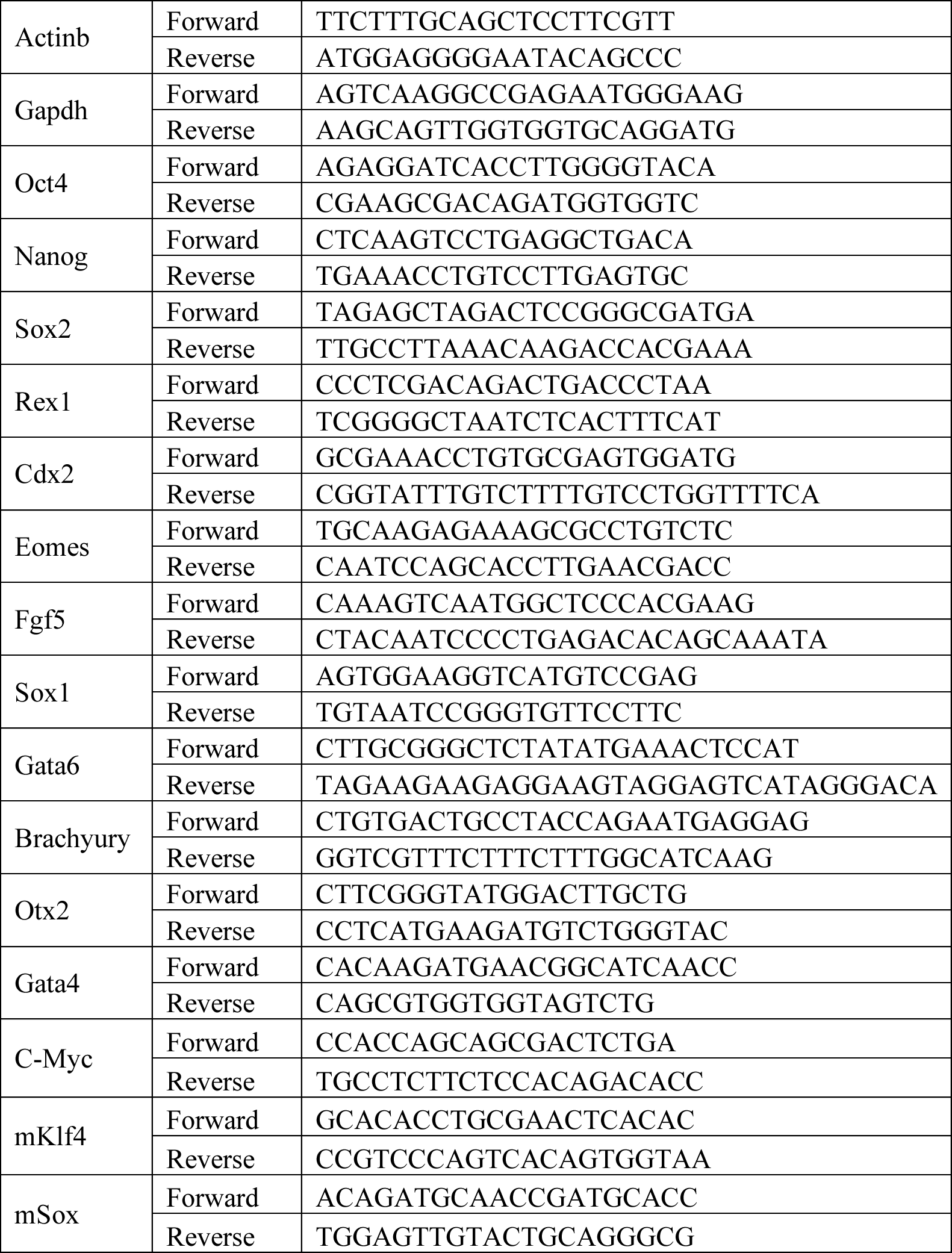

### Alkaline phosphatase (AP) staining

Alkaline phosphatase (AP) staining was performed with AP kit (Millipore SCR004) according to manufacturer protocol. Briefly, cells were fixated using 4% PFA for 2 minutes, and later washed with TBST. The reagents were then added to the wells, followed by an incubation of 10 minutes in RT. Stained plates were scanned, and positive AP+ colonies were counted to evaluate reprogramming efficiency at differential conditions.

### Apoptosis detection by Annexin-PI stain

Annexin-PI staining was performed with ‘The dead cell Apoptosis kit with Annexin V Alexa Fluor 488 and Propidium Iodide’ (ThermoScientific V13245), according to the manufacturer protocol. Examined cells were stained in both Annexin and PI in order to evaluate the rate of apoptotic and dead cells in the population. The cells were then analyzed by FACS (BD FACS ARIA III) in order to detect cell mortality and apoptosis rates.

### Co-immunoprecipitation (co-IP) and proteomics analysis

When used for overexpression experiments over-expression, HEK293T were transfected with our plasmids using JetPEI Transfection reagent (Polyplus). The cells were harvested after 48 hours, and protein interactions were examined by co-immunoprecipitation. Flag Coimmunoprecipitation experiments were performed using Flag magnetic beads (M2, Sigma M8823). Fresh protein extracts were incubated with the magnetic beads for 4 hours in 4 C degrees, in rotation. The beads were washed from the unbound proteins, and then incubated with elution buffer. The buffer contained 150mM Tris-Hcl (pH=7.4), 1.5 mM MgCl2, 150 mM NaCl, 0.5mg\ml 3Xflag peptide (sigma, F4799). HA co-immunoprecipitation was done with the same lysis buffer, but with Pierce anti-HA magnetic beads (ThermoScientific #88836). Elution was done with HA- peptide (Sigma I2149). Untagged and endogenous proteins co-immunoprecipitation was carried using protein-G Dynabeads (ThermoScientific); beads were incubated with the lysate for 4 hours in 4 C degrees, and elution was done by heat. The elution was analyzed by western blot analysis as detailed above or by mass-spectrometry analysis.

### Mass spectrometry sample preparation and analysis

The samples were then subjected to tryptic digestion, a process constituted from reduction, alkylation and finally digestion with trypsin (16 hours, 37 °C, at 1:50 trypsin-protein ration). Digestion was stopped, and the samples were desalted using solid-phase extraction columns. The samples were then subjected to liquid chromatography coupled to high resolution, tandem mass spectrometry (LC-MS/MS). The mobile phase used was H_2_O/Acetonitrile +0.1% formic acid, with the C18 column (reverse phase column). The peptide separation was performed using T3 HSS nano-column. Peptides were analyzed by a quadrupole orbitrap mass spectrometer, using flexion nanospray apparatus. Data was acquired in data dependent acquisition (DDA) mode, using a Top12 method. MS1 resolution was set to 60,000 (at 400m/z) and maximum injection time was set to 120msec. MS2 resolution was set to 17,500 and maximum injection time of 60 msec. Raw data was then imported to the Expressionist software (Genedata), and after peak detection, the peak list was searched using the Mascot algorithm as described in (Shalit et al., 2015). Data was normalized based on total ion current. Protein abundance was obtained by summing the three most intense, unique peptides per protein. Minimal cut-off for protein inclusion was ‘Ratio Sample/Control=3’, at least 2 unique peptides, and identification in at least 2 out of 3 repeats.

### Cell fractioning and Chromatin Extraction

The protocol for fractioning of proteins to Nucleoplasm, Cytoplasm and chromatin fraction was previously described (Huang et al., 2009; Toiber et al., 2013). Briefly, cells were resuspended in lysis buffer (10 mM HEPES pH 7.4, 10 mM KCl, 0.05% NP-40) and incubated 20 minutes on ice. The lysate was then centrifuged (14,000 RPM, 10 minutes), resulting in the separation of the nucleoplasm and the cytoplasmic fractions. Next, the pellet (containing the nuclei) is re-suspended in Low-salt Buffer (10 mM Tris-HCl pH 7.4, 0.2 mM MgCl2, 1% Triton-X 100) and incubated 15 minutes on ice. Subsequent centrifuge results in the separation of Chromatin and Nucleoplasm. The pellet (containing chromatin fraction) was then resuspended in HCl 0.2N and incubated on ice, then centrifuged. Supernatant was then kept and neutralized with Tris-HCl pH=8 1M (1:1 ratio). All buffers were supplemented with supplemented with 20mM of NEthylmaleimide (NEM - Sigma Aldrich E3876) and protease inhibitor cocktail (Sigma-Aldrich, P8340).

### SUMOylation inhibition and analysis

ES cells were treated with different concentration of Ubc9-inhibitor 2-D08 (Sigma-Aldrich SML1052) for 48 hours. Cells were then harvested and treated with NP-40 based lysis buffer, supplemented with 20mM of N-Ethylmaleimide (NEM- Sigma E3876) and protease inhibitor cocktail (Sigma P8340).

### Immunostaining of cells in culture

Cells subjected to immunostaining were washed three times with PBS and fixed with 4% paraformaldehyde for 10 minutes at room temperature. Cells were then permeabilized and blocked in 0.1% Triton, 0.1% Tween, and 5% FBS in PBS for 15 min at room temperature. Primary antibodies were incubated for two hours at room temperature and then washed with 0.1% Tween and 1% FBS in PBS three times. Next, cells were incubated with secondary antibody for one hour at room temperature, washed and counterstained with DAPI, mounted with Shandon Immu-Mount (ThermoScientific) and imaged. All secondary antibodies were diluted 1:200.

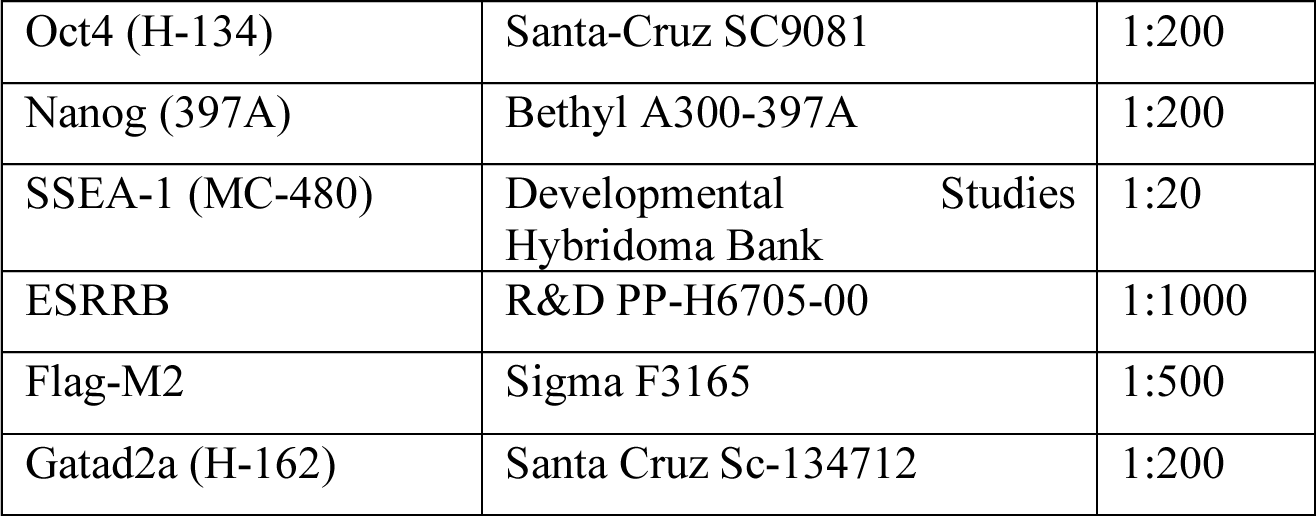

### Teratoma assay

iPSCs or ESCs grown in N2B27 2i/LIF cell lines were expended for over 8 passages and injected subcutaneously to the flanks of immune deficient NSG mice. After 4-6 weeks, all injected mice were sacrificed and the tumor mass extracted and fixed in 4% paraformaldehyde over-night. Slides were prepared from the paraffin embedded fixed tissue, which were next Hematoxylin … Eosin stained and inspected for representation of all three germ layers and confirmed by a pathology expert.

### Mouse embryo micromanipulation

Pluripotent ESCs/iPSCs were injected into BDF2 diploid blastocysts, harvested from hormone primed BDF1 6-week-old females. Microinjection into BDF2 E3.5 blastocysts placed in M2 medium under mineral oil was done by a flat-tip microinjection pipette. A controlled number of 10-12 cells were injected into the blastocyst cavity. After injection, blastocysts were returned to KSOM media (Zenith) and placed at 37^o^C until transferred to recipient females. Ten to fifteen injected blastocysts were transferred to each uterine horn of 2.5 days post coitum pseudo-pregnant females. All animal experiments described herein were approved by relevant Weizmann Institute IACUC (#00330111).

### PolyA-RNA-seq library preparation and analysis

Total RNA was isolated from indicated cell lines and extracted from Trizol pellets by Direct-zol RNA MiniPrep kit (Zymo), then utilized for RNA-Seq by TruSeq RNA Sample Preparation Kit v2 (Illumina) according to manufacturer’s instruction. RNA-seq data are deposited under GEO no. GSE102518. Tophat software version 2.0.10 was used to align reads to mouse mm10 reference genome (UCSC, December 2011). FPKM values were calculated over all genes in mm10 assembly GTF (UCSC, December 2011), using cufflinks (version 2.2.1). Genes with minimal expressed (FPKM>1) in at least one time point in each of the systems (Gatad2aKO/WT/WT*) were selected for clustering and PCA, resulting in 9,756 active genes. Unit normalized FPKM was calculated using the following formula R^i^_j_^*^ = R^i^_j_ / [max_j_(R^i^_j_)+1] where j is the sample index, i is the gene index and FPKM=1 is the transcription noise threshold, and max_j_(R^i^_j_) is the maximal level in each dataset. Hierarchical clustering was carried out using R pheatmap command. PCA analysis was carried out over the same set of 9,756 genes in unit normalization by R prcomp command.

### ATAC-seq library preparation and analysis

Cells were trypsinized and counted, 50,000 cells were centrifuged at 500*g* for 3 min, followed by a wash using 50 μl of cold PBS and centrifugation at 500*g* for 3 min. Cells were lysed using cold lysis buffer (10 mM Tris-HCl, pH 7.4, 10 mM NaCl, 3 mM MgCl_2_ and 0.1% IGEPAL CA-630). Immediately after lysis, nuclei were spun at 500*g* for 10 min using a refrigerated centrifuge. Next, the pellet was resuspended in the transposase reaction mix (25 μl 2× TD buffer, 2.5 μl transposase (Illumina) and 22.5 μl nuclease-free water). The transposition reaction was carried out for 30 min at 37 °C and immediately put on ice. Directly afterwards, the sample was purified using a Qiagen MinElute kit. Following purification, the library fragments were amplified using custom Nextera PCR primers 1 and 2 for a total of 12 cycles. Following PCR amplification, the libraries were purified using a *Qiagen*MinElute Kit and sequenced.

For ATAC-seq analysis, Reads were aligned to mm10 mouse genome using Bowtie2 with the parameter -X2000 (allowing fragments up to 2 kb to align). Duplicated aligned reads were removed using Picard MarkDuplicates tool with the command REMOVE_DUPLICATES=true. To identify chromatin accessibility signal we considered only short reads ((100bp) that correspond to nucleosome free region. To detect and separate accessible loci in each sample, we used MACS version 1.4.2-1 with --call-subpeaks flag (PeakSplitter version 1.0). Next, summits in previously annotated spurious regions were filtered out using a custom blacklist targeted at mitochondrial homologues. To develop this blacklist, we generated 10,000,000 synthetic 34mer reads derived from the mitochondrial genome. After mapping and peak calling of these synthetic reads we found 28 high-signal peaks for the mm10 genome. For all subsequent analysis, we discarded peaks falling within these regions.

### Enhancer Identification

H3K27ac peaks were detected using MACS version 1.4.2-1 and merged for all time points using bedtools merge command. All ATAC peaks were filtered to include only peaks which co-localized with the merged H3K27ac peaks, meaning only ATAC peaks that have H3K27ac mark on at least one of the time points were passed to further processing. Finally, the peaks from all samples were unified and merged (using bedtools unionbedg and merge commands), further filtered to reject peaks that co-localized with promoter or exon regions based on mm10 assembly (UCSC, December 2011). Finally, we were left with 52,473 genomic intervals which we annotated as active enhancers.

Enhancers were considered as differential if both their ATAC-seq and H3K27ac signals show significant change during reprogramming (min z-score<0.5, max z-score>1.5, for both chromatin marks). 12,153 enhancers were found to be differential and were used for correlation analysis **(Fig. 4C)**. Z-scores were calculated as following: Shortly, the genomic interval is divided to 50bp size bins, and the coverage in each bin is estimated. Each bin is then converted to z-score by normalizing each position by the mean and standard deviation of the sample noise (X^j=(Xj-μnoise)/σnoise). Noise parameters were estimated for each sample from 6*10^7^ random bp across the genome. Finally, the 3^rd^ highest bin z-score of each interval is set to represent the coverage of that interval. ATAC-seq data are deposited under GEO no. GSE102518.

### Whole-Genome Bisulfite Sequencing (WGBS) Library preparation and analysis

DNA was isolated from snap-frozen cells using the Quick-gDNA mini prep kit (Zymo). DNA was then converted by bisulfite using the EZ DNA Methylation-Gold kit (Zymo). Sequencing libraries were created using the EpiGenome Methyl-Seq (Epicenter) and sequenced. The sequencing reads were aligned to the mouse mm10 reference genome (UCSC, December 2011), using a proprietary script based on Bowtie2. In cases where the two reads were not aligned in a concordant manner, the reads were discarded. Methylation levels of CpGs calculated by RRBS and WGBS were unified. Mean methylation was calculated for each CpG that was covered by at least 5 distinct reads (X5). Average methylation level in various genomic intervals was calculating by taking the average over all covered X5 covered CpG sites in that interval. WGBS data are available to download from NCBI GEO, under super-series GSE102518.

### ChIP-seq library preparation and analysis

Cells were crosslinked in formaldehyde (1% final concentration, 10 min at room temperature), and then quenched with glycine (5 min at room temperatureσ. Fixated cells were then lysed in 50 mM HEPES KOH pH…7.5, 140 mM NaCl, 1 mM EDTA, 10% glycerol, 0.5% NP-40 alternative, 0.25% Triton supplemented with protease inhibitor at 4 °C (Roche, 04693159001), centrifuged at 950*g* for 10 min and re-suspended in 0.2% SDS, 10 mM EDTA, 140 mM NaCl and 10 mM Tris-HCl. Cells were then fragmented with a Branson Sonifier (model S-450D) at *4…°C to size ranges between 200 and 800 bp and precipitated by centrifugation. Antibody was pre-bound by incubating with Protein-G Dynabeads (Invitrogen 100-07D) in blocking buffer (PBS supplemented with 0.5% TWEEN and 0.5% BSA) for 1 h at room temperature. Washed beads were added to the chromatin lysate for incubation as detailed in the table below. Samples were washed five times with RIPA buffer, twice with RIPA buffer supplemented with 500 mM NaCl, twice with LiCl buffer (10 mM TE, 250mM LiCl, 0.5% NP-40, 0.5% DOC), once with TE (10Mm Tris-HCl pH 8.0, 1mM EDTA), and then eluted in 0.5% SDS, 300…mM NaCl, 5 mM EDTA, 10 mM Tris HCl pH…8.0. Eluate was incubated treated sequentially with RNaseA (Roche, 11119915001) for 30 min and proteinase K (NEB, P8102S) for 2 h in 65 °C for 8 h, and then. DNA was purified with The Agencourt AMPure XP system (Beckman Coulter Genomics, A63881). Libraries of cross-reversed ChIP DNA samples were prepared according to a modified version of the Illumina Genomic DNA protocol, as described previously. ChIP-seq data are available to download from NCBI GEO, under super-series GSE102518.

Alignment and peak detection: We used bowtie2 software to align reads to mouse mm10 reference genome (UCSC, December 2011), with default parameters. We identified enriched intervals of all measured proteins using MACS version 1.4.2-1. We used sequencing of whole-cell extract as control to define a background model. Duplicate reads aligned to the exact same location are excluded by MACS default configuration.

Chromatin modification profile estimation in TSS, TES and in enhancers: H3K27ac modification coverage in the genomic intervals was calculated using in-house script. Shortly, the genomic interval is divided to 50bp size bins, and the coverage in each bin is estimated. Each bin is then converted to z-score by normalizing by the mean and standard deviation of the sample noise (X^j=(Xj-μnoise)/σnoise). Noise parameters were estimated for each sample from 6*10^7^ random bp across the genome. Finally, the 3^rd^ highest bin z-score of each interval is set to represent the coverage of that interval.

|  |  |  |
| --- | --- | --- |
| H3K27ac | Abcam ab4729 | 6H |
| Oct4 | Santa Cruz SC8628 | Overnight |
| Klf4 | R&D AF3158 | Overnight |
| Sox2 | Millipore AB5603 | Overnight |
| C-Myc | Santa Cruz sc764 | Overnight |
| Chd4 | Abcam ab70469 | Overnight |
| Mbd3 | Bethyl laboratories A302-528\9 | Overnight |

### Microscopy live image acquisition and analysis

Isogenic secondary OKSM inducible Gatad2a-KO and Gatad2a-WT secondary MEFs harboring ΔPE-Oct4-GFP pluripotency reporter and constitutively expressed nuclear mCherry marker (**Fig. S2A**), were plated in 24-well plates at low densities (˜120 cells per well). Reprogramming was than imaged (beginning 72 hours after DOX administration) using AxioObserver Z1 (Zeiss) in %5 O2, %5 CO2, 37C controlled conditions. Plates were taken out at day 3.5 for media replacement (without passaging/splitting) and put back for the automated live imaging stage. Full well mosaic images were taken every 12 hours for 4 days (days 3-7 of reprogramming) at 5x magnification, including phase contrast and two fluorescent wavelength images.

### Data Availability

All RNA-seq, ATAC-seq, ChIP-seq and WGBS methylation data are available to download from NCBI GEO, under super-series GSE102518. They will be publically released upon printed publication of this study after peer review or following special requests.

